# Optogenetic actuator/ERK biosensor circuits identify MAPK network nodes that shape ERK dynamics

**DOI:** 10.1101/2021.07.27.453955

**Authors:** Coralie Dessauges, Jan Mikelson, Maciej Dobrzyński, Marc-Antoine Jacques, Agne Frismantiene, Paolo Armando Gagliardi, Mustafa Khammash, Olivier Pertz

**Affiliations:** Institute of Cell Biology, University of Bern, Baltzerstrasse 4, 3012 Bern, Switzerland; Department of Biosystems Science and Engineering, ETH Zurich, Mattenstrasse 26, 4058 Basel, Switzerland

**Keywords:** ERK dynamics, MAPK network, signaling robustness, optogenetics, single cell biology

## Abstract

Combining single-cell measurements of ERK activity dynamics with perturbations provides insights into the MAPK network topology. We built circuits consisting of an optogenetic actuator to activate MAPK signaling and an ERK biosensor to measure single-cell ERK dynamics. This allowed us to conduct RNAi screens to investigate the role of 50 MAPK proteins in ERK dynamics. We found that the MAPK network is robust against most node perturbations. We observed that the ERK-RAF and the ERK-RSK2-SOS negative feedbacks operate simultaneously to regulate ERK dynamics. Bypassing the RSK2-mediated feedback, either by direct optogenetic activation of RAS, or by RSK2 perturbation, sensitized ERK dynamics to further perturbations. Similarly, targeting this feedback in a human ErbB2-dependent oncogenic signaling model increased the efficiency of a MEK inhibitor. The RSK2-mediated feedback is thus important for the ability of the MAPK network to produce consistent ERK outputs and its perturbation can enhance the efficiency of MAPK inhibitors.

## Introduction

The extracellular signal-regulated kinase (ERK) is part of the mitogen-activated protein kinase (MAPK) signaling network and regulates a large variety of fate decisions. While ERK can be activated by several extracellular inputs, ERK signaling has mostly been studied in the context of receptor tyrosine kinases (RTKs). Upon binding of their cognate growth factors (GFs), RTKs activate a complex signaling cascade with the following hierarchy: (1) recruitment of adaptor molecules such as GRB2 (Schlessinger 2000), (2) activation of RAS GTPases through Guanine nucleotide exchange factors (GEFs) and GTPase activating proteins (GAPs) (Cherfils and Zeghouf 2013), (3) triggering of a tripartite RAF, MEK, ERK kinase cascade that is further regulated by a variety of binding proteins (Lavoie et al. 2020), (4) ERK-mediated phosphorylation of a large number of substrates. Due to its central role in fate decisions, MAPK network dysregulation is causative for a large number of diseases including cancer (Rauen 2013; Samatar and Poulikakos 2014).

As for other signaling pathways (Purvis and Lahav 2013), temporal patterns of ERK activity, hereafter referred to as ERK dynamics, rather than steady states control fate decisions (Santos et al. 2007; Avraham and Yarden 2011; Ryu et al. 2015; Albeck et al. 2013). These specific ERK dynamics have been shown to arise from feedbacks in the MAPK network. For example, a negative feedback (NFB) from ERK to RAF can produce adaptive or oscillatory ERK dynamics (Santos et al. 2007; Kholodenko et al. 2010; Avraham and Yarden 2011). The ERK-RAF NFB was also shown to buffer against MAPK node perturbations (Sturm et al. 2010; Fritsche-Guenther et al. 2011). This property might allow cells to produce consistent ERK outputs despite heterogeneous node expression (Blüthgen and Legewie 2013). In this work, we specifically refer to the ability of the MAPK network to produce consistent ERK dynamics in presence of node perturbations as signaling robustness. While several NFBs have been mapped experimentally in the MAPK network (Lake et al. 2016), their contribution to this signaling robustness and shaping ERK dynamics remains largely unknown.

Single-cell biosensor imaging has provided new insights into MAPK signaling that were not accessible with biochemical, population-averaged measurements. This showed that the MAPK network can produce a wide variety of ERK dynamics such as transient (Ryu et al. 2015), pulsatile (Albeck et al. 2013), oscillatory (Shankaran et al. 2009) and sustained dynamics (Ryu et al. 2015; Blum et al. 2019). Mathematical modeling has provided insights into the network’s structures that decode different signaling inputs into specific ERK dynamics (Santos et al. 2007; Shankaran et al. 2009; Nakakuki et al. 2010; Ryu et al. 2015). Combined modeling/experimental approaches helped to shed light on various subparts of the MAPK network, including the epidermal growth factor receptor (EGFR) module (Koseska and Bastiaens 2020), the RAS module (Schmick et al. 2015; Erickson et al. 2019), and the tripartite RAF/MEK/ERK cascade (Ferrell and Bhatt 1997; Kholodenko 2000; Orton et al. 2005; Santos et al. 2007; Ryu et al. 2015; Kochańczyk et al. 2017; Arkun and Yasemi 2018). However, the low experimental throughput to measure ERK dynamics, or other MAPK network nodes, has precluded a global understanding of the specific functions of the nodes present in the network.

Here, we built multiple genetic circuits consisting of optogenetic actuators together with an ERK biosensor to simultaneously activate ERK from different nodes in the MAPK network and report single-cell ERK dynamics. These circuits allowed us to investigate the role of 50 MAPK signaling nodes in ERK dynamics regulations with RNA interference (RNAi). We observed that most perturbations of individual nodes resulted in mild ERK dynamics phenotypes despite targeting major MAPK signaling nodes. Further, the ERK dynamics induced by various perturbations suggest that two NFBs (ERK-RAF and ERK-RSK2-SOS) act simultaneously to regulate ERK dynamics. Targeting the RSK2-mediated NFB increased the efficiency of additional MAPK network perturbations both in our optogenetic systems and in an ErbB2-driven oncogenic ERK signaling model. This suggests that the RSK2-mediated feedback plays a role in MAPK signaling robustness and can be targeted for potent inhibition of oncogenic ERK signaling.

## Results

### An optogenetic actuator-biosensor genetic circuit to study input-dependent ERK dynamics

In order to measure ERK dynamics in response to dynamic RTK input, we built a genetically-encoded circuit made of an optogenetic RTK actuator and an ERK biosensor (Figure 1A). We chose optoFGFR, which consists of a myristoylated intracellular domain of the fibroblast growth factor receptor 1 (FGFR1) fused to a CRY2 domain and tagged with mCitrine (Kim et al. 2014). Upon stimulation with blue light, optoFGFR dimerizes and trans*-*autophosphorylates, leading to the activation of the MAPK/ERK, phosphoinositide 3-kinase (PI3K)/AKT, and phospholipase C (PLC)/Ca^2+^ pathways. As ERK biosensor, we used ERK-KTR-mRuby2 that is spectrally compatible with optoFGFR. ERK-KTR reversibly translocates from the nucleus to the cytosol upon ERK activation (Regot et al. 2014). We used a nuclear Histone 2B (H2B)-miRFP703 marker to identify and track single cells. After stably inserting these constructs into murine NIH3T3 fibroblasts, we used automated time-lapse microscopy to stimulate selected fields of view with defined blue light input patterns to activate optoFGFR. The corresponding ERK-KTR/H2B signals were recorded with a 1-minute temporal resolution. We observed that a 100 ms light pulse leads to reversible ERK-KTR translocation from the nucleus to the cytosol, indicative of transient ERK activation (Figure 1B, Appendix Movie S1). At the end of each experiment, we imaged the mCitrine signal to evaluate optoFGFR expression levels. We built a computer vision pipeline to automatically track each nucleus, compute ERK activity as the cytosolic/nuclear ratio of the ERK-KTR signals and correlate single-cell ERK responses with optoFGFR levels (Figure 1C). We then use this pipeline to evaluate the sensitivity and specificity of our system with dose response experiments using the FGFR inhibitor SU5402, the RAF inhibitor RAF709, the MEK inhibitor U0126 and the ERK inhibitor SCH772984 (Appendix Figure S1A).

**Figure 1:**
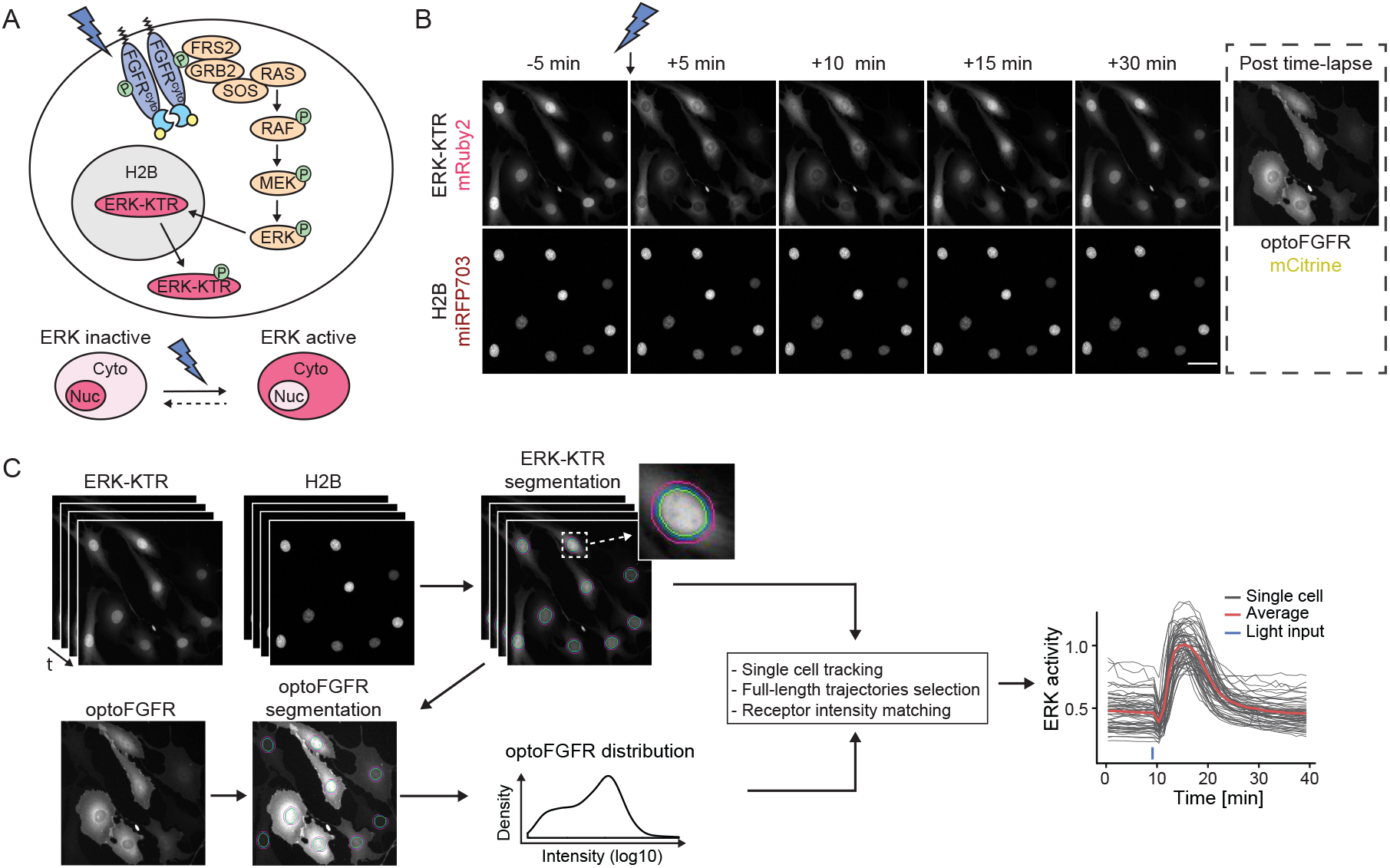
An optogenetic actuator-biosensor genetic circuit to study input-dependent ERK dynamics. **(A)** Schematic representation of the optoFGFR system consisting of the optogenetic FGF receptor (optoFGFR) tagged with mCitrine, the ERK biosensor (ERK-KTR) tagged with mRuby2 and a nuclear marker (H2B) tagged with miRFP703. **(B)** Time lapse micrographs of ERK-KTR dynamics in response to a 470 nm light pulse. Using a 20x air objective, ERK-KTR and H2B channels were acquired every 1 minute and the optoFGFR channel was acquired once at the end of the experiment. Scale bar: 50 μm. **(C)** Image analysis pipeline developed to quantify single-cell ERK dynamics. Nuclear and cytosolic ERK-KTR signals were segmented based on the H2B nuclear mask. Single-cell ERK activity was then calculated as the cytosolic/nuclear ERK-KTR ratio. Single-cell optoFGFR intensity was measured under the cytosolic ERK-KTR mask and used as a proxy for single-cell optoFGFR expression.

To evaluate light-dependent optoFGFR activation dynamics, we engineered a mScarlet-tagged optoFGFR that is spectrally orthogonal to CRY2 absorption (Appendix Figure S1B). Total internal reflection (TIRF) microscopy visualized the formation of optoFGFR clusters in response to blue light-mediated dimerization in the plasma membrane (Appendix Figure S1B, blue arrows, Appendix Movie S2). Consistently with CRY2’s dissociation half-life (Duan et al. 2017), these optoFGFR clusters appeared within 20 seconds after a blue light pulse and disappeared after ∼ 5 minutes (Appendix Figure S1C). We assume that optoFGFR is active in its clustered form in which transphosphorylation occurs and inactive in its monomeric form due to tonic cytosolic phosphatase activity (Lemmon et al. 2016). As documented previously (Kim et al. 2014), light stimulation also triggered optoFGFR endocytosis (Appendix Figure S1B, red arrows).

Directly following light stimulation, we systematically observed a short ERK inactivation period, that we refer to as “dip”, lasting 2-3 minutes before activation of a strong ERK activity (Appendix Figure S1D, green rectangle). This light-induced ERK dip was insensitive to SCH772984-mediated ERK inhibition but could be suppressed by Cyclosporin A-mediated calcineurin inhibition. Calcineurin is a Ca^2+^-dependent phosphatase that dephosphorylates Ser383 in Elk1 (Sugimoto et al. 1997). As ERK-KTR contains an Elk-1 docking domain phosphorylated by ERK (Regot et al. 2014), we hypothesized that it could be negatively affected by optoFGFR-evoked Ca^2+^ input (Kim et al. 2014) (Appendix Figure S1E). Consistently, Ionomycin-evoked increase in cytosolic Ca^2+^ induced a dip in absence of light stimulation (Appendix Figure S1F).

### Different optoFGFR inputs trigger transient, oscillatory and sustained ERK dynamics

Next, we characterized optoFGFR-triggered ERK dynamics in response to a single light pulse of different intensities and durations (Figure 2A). As ERK dynamics depended on light power density, as well as pulse duration, we defined the light dose (D, mJ/cm^2^) as their product to quantify the total energy received per illuminated area. To characterize ERK dynamics, we extracted the amplitude at the maximum of the peak (maxPeak), and the full width at half maximum (FWHM) of the ERK trajectories (Figure 2B). With increasing light doses, ERK peaks increased both in duration and amplitude, until the latter reached saturation. Based on these observations, we selected 180 mW/cm^2^ and 100 ms (D = 18 mJ/cm^2^) as the minimal light input to generate an ERK transient of maximal amplitude. Using this light dose, we then investigated ERK dynamics in response to multiple light pulses delivered at different intervals (Figure 2C). All stimulation regimes led to identical maximal ERK amplitude (Figure EV1A) and adaptation kinetics when optoFGFR input ceased (Figure EV1B). Repeated light inputs applied at 10- or 20-minute intervals evoked population-synchronous ERK transients. In contrast, repeated light inputs applied at higher frequencies (2-minute intervals) led to sustained ERK dynamics. Given CRY2’s 5-minute dissociation half-life (Appendix Figure S1B-C) (Duan et al. 2017), this suggests that light pulses delivered at a 2-minute interval reactivate optoFGFR faster than it deactivates, leading to sustained optoFGFR activity. Hierarchical clustering of ERK responses to sustained optoFGFR input highlighted the presence of sustained and oscillatory single-cell ERK dynamics (Figure 2D). Classification of ERK trajectories based on optoFGFR expression revealed that sustained/oscillatory ERK dynamics correlated with high/low optoFGFR levels (Figure 2E, Appendix Movie S3). Oscillatory ERK dynamics were also observed in optoFGFR high expressing cells in response to low light input (Figure 2F). Thus, sustained optoFGFR input can trigger sustained or oscillatory ERK dynamics depending on the input strength, a combination of light energy and optoFGFR expression.

**Figure 2:**
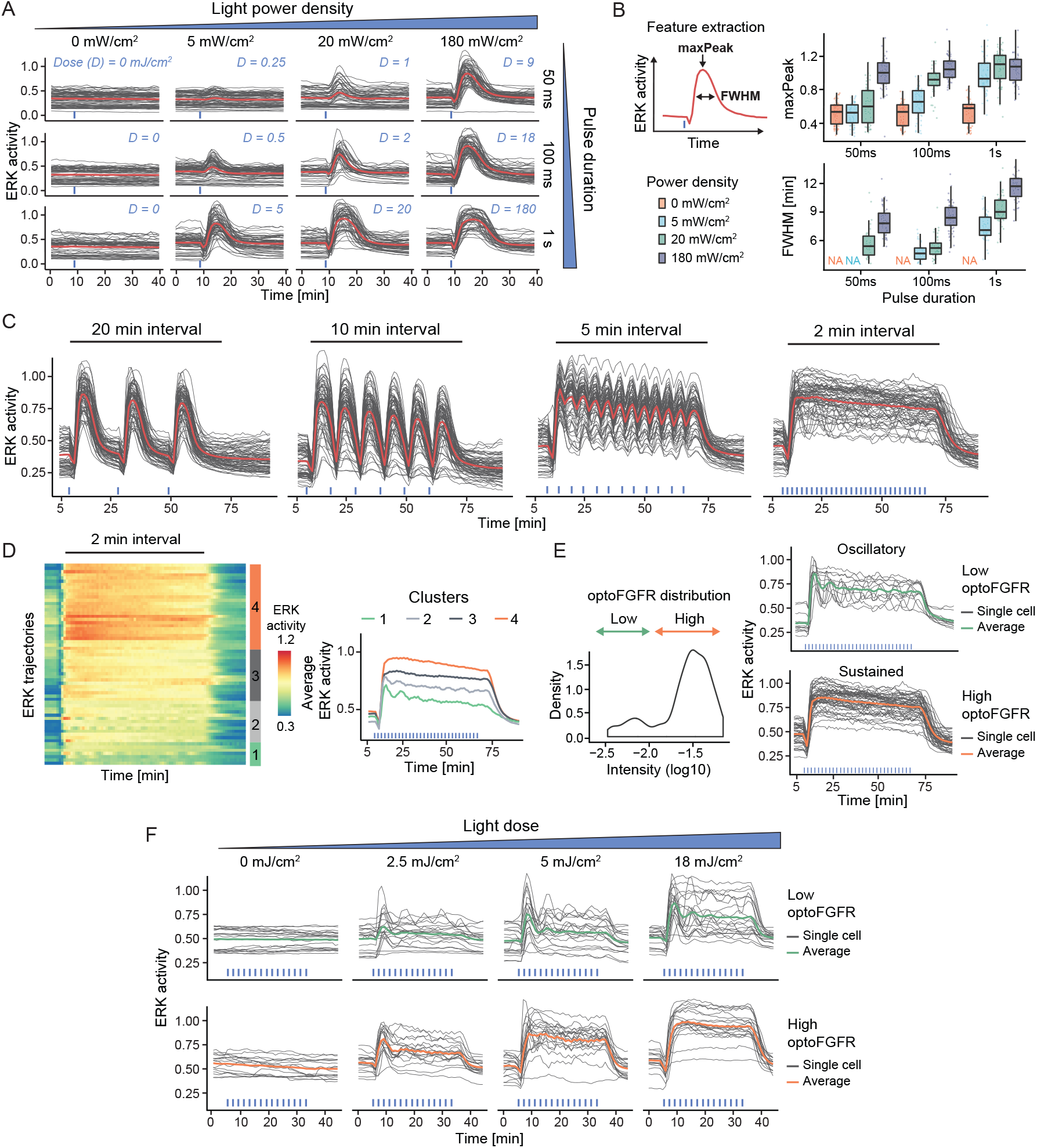
Different optoFGFR inputs trigger transient, oscillatory and sustained ERK dynamics. **(A)** ERK responses to increasing light power densities and pulse durations of 470 nm transient light input. The light dose “D” is calculated as the product of the power density and pulse duration. **(B)** Quantification of the maxPeak (maximal ERK amplitude of the trajectory) and the FWHM (full width at half maximum) of single-cell ERK responses shown in (A) (N_min_ = 40 cells per condition). **(C)** ERK responses to 470 nm light pulses delivered every 20, 10, 5 and 2 minutes respectively (D = 18 mJ/cm^2^). **(D)** Hierarchical clustering (Euclidean distance and Ward D2 linkage) of trajectories from the 2-minute interval stimulation shown in (C) (referred to as “sustained”) (N = 60 cells). The number of clusters was empirically defined to resolve the different ERK dynamics. The average ERK responses per cluster are displayed on the right. **(E)** Separation of the trajectories shown in (D) in low and high optoFGFR cells, based on the log10 intensity of optoFGFR-mCitrine. **(F)** ERK responses to increasing doses of sustained optoFGFR input. Single-cell ERK trajectories were divided in low (top panel) and high (bottom panel) optoFGFR expression.

### ERK dynamics evoked by optoFGFR versus endogenous RTKs highlight different MAPK regulatory mechanisms

Because of the absence of an ectodomain, optoFGFR must be considered as a prototypic RTK that lacks some regulatory mechanisms inherent to the native FGFR. To evaluate if optoFGFR is relevant for studying the MAPK network, we compared ERK dynamics evoked by optoFGFR inputs versus stimulation of the endogenous FGFR or EGFR using increasing concentrations of basic FGF (bFGF) and EGF. All bFGF concentrations led to an ERK peak similar in amplitude to sustained optoFGFR input (Figure 3A, EV1C, compared to EV1A). However, FGFR inputs led to different ERK dynamics than optoFGFR: 1 ng/ml bFGF led to damped ERK oscillations followed by steady state sustained ERK activity, while 10 and 100 ng/ml bFGF concentrations led to a first ERK peak followed by a strong adaptation. The biphasic behavior induced by increasing bFGF concentrations was previously documented to emerge from the competition of bFGF for FGFR and heparan sulfate proteoglycan co-receptors (Kanodia et al. 2014; Blum et al. 2019). It is thus not surprising that optoFGFR, that lacks these extracellular interactions, produced different ERK dynamics than FGFR. All EGF concentrations led to an ERK peak similar in amplitude to optoFGFR and FGFR inputs (Figure 3B, EV1D). As for bFGF, 1 ng/ml EGF concentration evoked damped oscillatory ERK dynamics that decreased at higher EGF concentrations. However, EGFR inputs led to strong ERK adaptation, not observed in response to optoFGFR inputs, suggesting the existence of different regulatory mechanisms.

**Figure 3:**
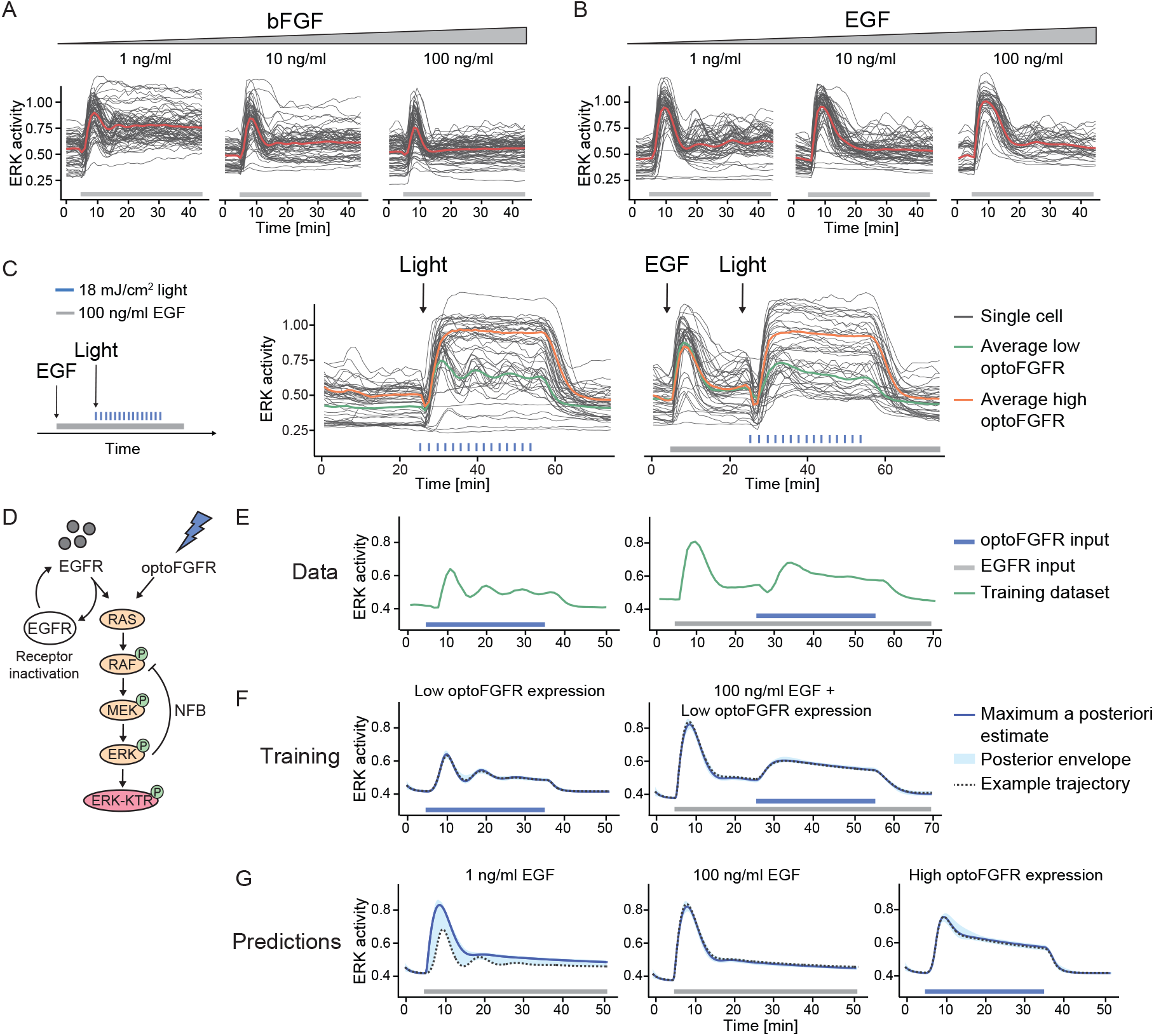
ERK dynamics evoked by optoFGFR versus endogenous RTKs highlight different MAPK regulatory mechanisms. **(A-B)** Single-cell ERK trajectories under increasing concentrations of sustained **(A)** bFGF or **(B)** EGF input added at t = 5 minutes. **(C)** ERK responses of cells stimulated with sustained optoFGFR input (D = 18 mJ/cm^2^) at t = 24 minutes without or with 100 ng/ml EGF sustained pre-stimulation at t = 5 minutes. Average ERK responses for optoFGFR high and low expression levels are shown (N = 20 cells for low and high optoFGFR, randomly selected out of at least 80 cells). **(D)** Mathematical model topology consisting of the RAS GTPase, the MAPK three-tiered (RAF, MEK, ERK) network and the ERK-KTR reporter. EGFR and optoFGFR inputs both activate the RAS/RAF/MEK/ERK cascade and the ERK-RAF NFB. EGFR activity is under receptor-dependent regulations. **(E)** Training dataset consisting of the average ERK responses evoked by sustained low optoFGFR input with or without pre-stimulation with 100 ng/ml sustained EGF. **(F)** Simulation of ERK responses from the training dataset, including the maximum a posteriori (MAP) estimate, the posterior envelope indicating the predictive density of our estimation, as well as an example trajectory. **(G)** Predictions of the model for ERK responses evoked by 1 ng/ml EGF, 100 ng/ml EGF and sustained high optoFGFR inputs. Note that for low EGFR input (1 ng/ml), the model predicts both adaptive and oscillatory ERK responses.

Both oscillatory and transient ERK dynamics can be explained by the presence of NFB (Kholodenko et al. 2010). Thus, we wondered if the different ERK dynamics induced by optoFGFR or EGFR input emerge from differences in downstream NFBs. We reasoned that if EGFR induces different NFBs than optoFGFR, pre-stimulating cells with EGF should activate these feedbacks, and affect subsequent optoFGFR-evoked ERK dynamics. To test this, we pre-stimulated cells with sustained EGFR input, subsequently applied sustained optoFGFR input, and evaluated ERK dynamics (Figure 3C). Pre-stimulation with 100 ng/ml EGF led to the characteristic adaptive ERK transient. Subsequent application of optoFGFR input yielded sustained ERK responses similar in amplitude and duration to non-pre-stimulated cells. However, EGF pre-stimulation led to a reduction of synchronous optoFGFR-evoked ERK oscillations in low optoFGFR expressing cells.

To provide intuition about the MAPK network circuitries leading to different ERK dynamics in response to optoFGFR and EGFR inputs, as well as the origin of the oscillatory behavior, we built a mathematical model consisting of the RAS GTPase and the three-tiered RAF/MEK/ERK network (Figure 3D, Appendix Table S1). We used ordinary differential equations with Michaelis-Menten kinetics (see Material and methods, Appendix Table S2 and S3). To account for the oscillatory ERK dynamics in response to EGFR and optoFGFR inputs, we included the well-documented ERK-RAF NFB (Kholodenko et al. 2010; Santos et al. 2007; Fritsche-Guenther et al. 2011; Blum et al. 2019). We also included a receptor level inactivation process for EGFR, but not for optoFGFR, to account for EGF-dependent regulatory mechanisms. We used a Bayesian inference approach (Mikelson and Khammash 2020) to infer the model parameters from averaged ERK trajectories in response to sustained low optoFGFR input with or without sustained EGFR pre-stimulation (Figure 3E). After identification of parameters that allowed the model to capture the training dataset (Figure 3F), we simulated ERK dynamics evoked by low EGFR input (adaptative, oscillatory ERK dynamics), high EGFR input (adaptative ERK dynamics without oscillation) and sustained high optoFGFR input (sustained ERK dynamics) (Figure 3G). We observed that our model with a NFB and EGFR inactivation was able to predict ERK dynamics evoked by different EGFR and optoFGFR input strengths, while two simpler models (one with only the EGFR inactivation reaction, but no NFB (Figure EV1E-G) and one with only the NFB, but no EGFR inactivation (Figure EV1H-J) were not able to reproduce experimentally observed ERK dynamics.

This suggested that oscillatory optoFGFR-evoked ERK dynamics emerge from a NFB also present downstream of endogenous EGFR, while additional regulatory mechanisms seem to be required for the strong ERK transient adaptation following EGFR input. These mechanisms might consist of receptor-level regulations such as endocytosis, which was recently shown to be an important regulator of the transient adaptive EGF-triggered ERK dynamics in different cell systems (Kiyatkin et al. 2020; Gerosa et al. 2020). While optoFGFR also gets endocytosed (Appendix Figure S1B, (Kim et al. 2014)), it most likely is insensitive to inactivation by endosome acidification since it lacks an ectodomain (Huotari and Helenius 2011). Additionally, light-mediated optoFGFR dimerization might occur both at the plasma and endo-membranes, allowing for reactivation of endocytosed optoFGFR. The hypothesis that a receptor level mechanism is important for strong adaptation was further supported by inhibition of optoFGFR with the FGFR kinase inhibitor (SU5402), which shifted ERK dynamics from sustained to transient in a dose response-dependent manner (Figure EV1K). Thus, these results suggest that optoFGFR lacks receptor-dependent regulatory mechanisms but allows us to investigate the intracellular MAPK feedback structure shaping ERK dynamics. In our model, we used the well-established ERK-RAF NFB. However, several NFBs have been mapped in the MAPK signaling cascade, whose role in shaping ERK dynamics is still unknown and which could also be responsible for the observed oscillatory ERK dynamics.

### RNA interference screen reveals that ERK dynamics remain unaffected in response to perturbation of most MAPK signaling nodes

We then explored the network circuitry that shapes optoFGFR-evoked ERK dynamics with an RNA interference (RNAi) screen targeting 50 MAPK signaling nodes. We focused our screen on sustained optoFGFR input which captured the largest amount of information about ERK dynamics when compared to other stimulation schemes: it led to sustained and oscillatory ERK dynamics (Figure 2E,F) while recapitulating the rapid increase of ERK activity and adaptation observed with transient input (Figure EV1A,B). We used a bioinformatic approach to select 50 known interactors of the tripartite RAF/MEK/ERK cascade downstream of the FGFR receptor that were detected in a NIH3T3 proteome (Schwanhäusser et al. 2011) (Figure 4A, Appendix Table S4). We used the siPOOL technology to specifically knockdown (KD) these 50 MAPK signaling nodes while limiting off-target effects (Hannus et al. 2014). We first validated KD efficiency by quantifying transcript levels with different siPOOL concentrations targeting the ERK and MEK isoforms (Figure EV2A) and observed strong KD with 10 nM siRNA concentration. We then evaluated the effect of *ERK1* or *ERK2* KD on ERK dynamics. We observed only subtle phenotypes compared to the non-targeting siRNA (*CTRL*) used as negative control (Figure 4B), even though efficient KD was observed at protein level (Figure 4C). However, combined *ERK1/ERK2* KD strongly suppressed ERK dynamics indicating that the latter is not affected by the perturbation of individual ERK isoforms as previously reported (Fritsche-Guenther et al. 2011; Ornitz and Itoh 2015). Due to its strong phenotype, we used *ERK1/ERK2* KD as positive control throughout our screen.

**Figure 4:**
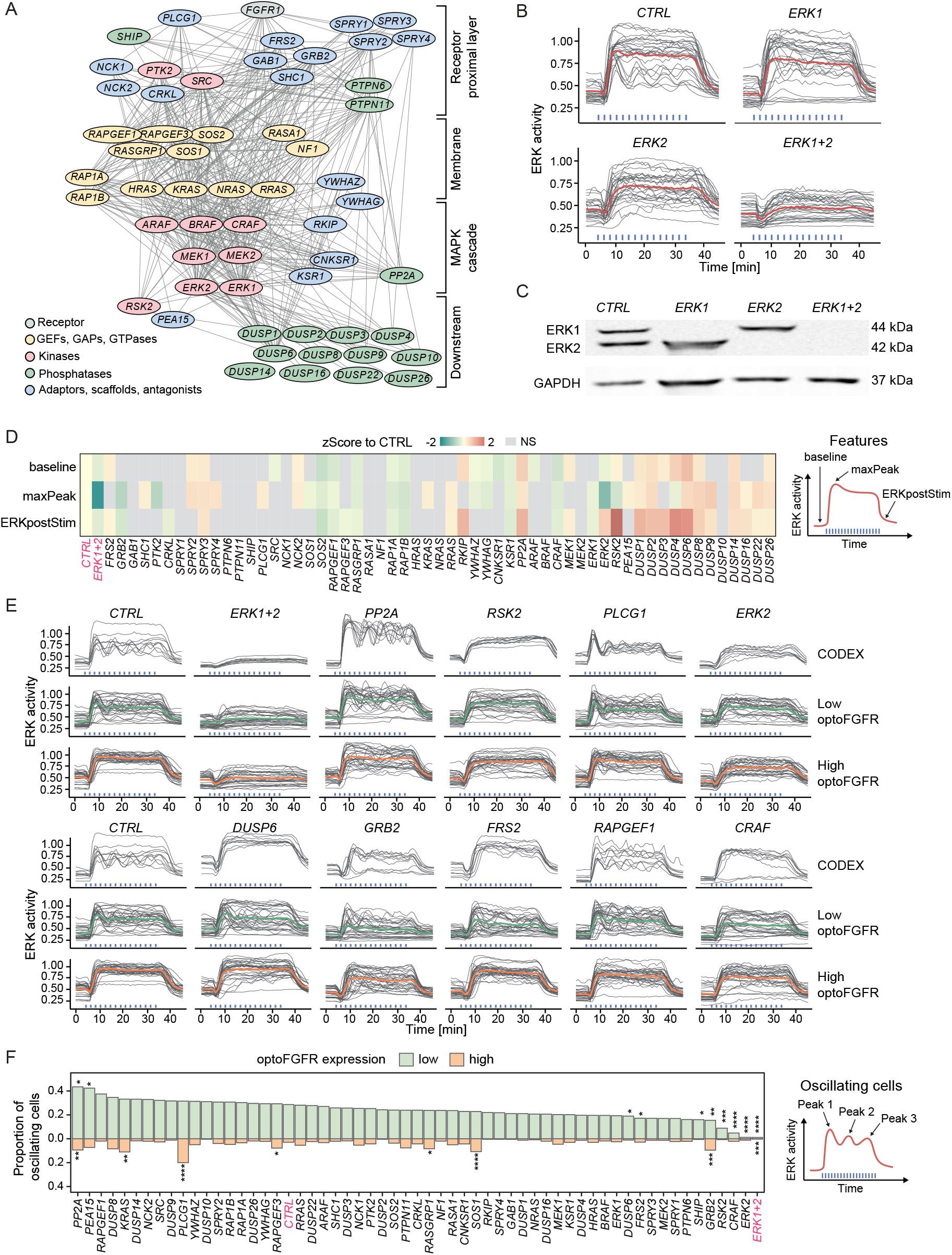
RNA interference screen reveals that ERK dynamics remain unaffected in response to perturbation of most MAPK signaling nodes. **(A)** RNAi perturbation targets referred to by their protein names. Nodes were spatially grouped based on the hierarchy of interactions within the MAPK network and color-coded for their function. **(B)** ERK responses to sustained optoFGFR input (D = 18 mJ/cm^2^) in cells transfected with 10 nM siRNA against *ERK1, ERK2* or a 5 nM combination of each (*ERK1+2*). A non-targeting siRNA (*CTRL*) was used as control (N = 15 cells from low and high optoFGFR levels). **(C)** Western blot analysis of cells transfected with 10 nM siRNA against *ERK1, ERK2* or a 5 nM combination of each (*ERK1+2*). **(D)** Z-Score evaluation of the baseline, maxPeak and ERKpostStim of single-cell ERK responses under sustained high optoFGFR input (D = 18 mJ/cm^2^). The z-score was calculated by comparing each RNAi perturbation to the *CTRL* KD (N_min_ = 126 cells per treatment, from 3 technical replicates). Non-significant (NS) results are in grey (see Figure EV3A for statistical results). **(E)** Single-cell ERK trajectories (sustained optoFGFR input, D = 18 mJ/cm^2^) for the RNAi perturbations classified with the highest accuracy by CODEX. Top lines show single-cell ERK trajectories for which CODEX had the highest classification confidence in the validation set (N = 10). Bottom lines show single-cell ERK trajectories for low and high optoFGFR cells (N = 30 for each condition, randomly selected out of at least 212 cells per perturbation from 3 technical replicates). For easier visualization, the CTRL condition is shown twice. **(F)** Proportion of oscillating cells (trajectories with at least 3 peaks) per RNAi perturbation for low and high optoFGFR expression (sustained optoFGFR input, D = 18 mJ/cm^2^, N_min_ = 61 cells for low and 126 for high optoFGFR per perturbation from 3 technical replicates). Perturbations were ordered based on the proportion of oscillating cells with low optoFGFR expression. Statistical analysis was done using a pairwise t-test, comparing each perturbation against the *CTRL* for each receptor level independently (*<0.05, **<0.005, ***<0.0005, ****<0.00005, FDR p-value correction method).

We performed three replicates of the screen targeting the 50 nodes. Despite efficient KD quantified for different nodes (Figure EV2B), visual inspection of ERK trajectories only revealed subtle ERK dynamics phenotypes for a limited number of node perturbations (Figure EV2C,D). We used a feature-based approach to evaluate the effect of each perturbation on ERK dynamics. We focused our analysis on ERK responses evoked by high optoFGFR input to limit the single-cell heterogeneity due to optoFGFR expression variability. We quantified the average ERK activity before stimulation (baseline), the maximal ERK amplitude during stimulation (maxPeak), and the ERK amplitude at a fixed time point after response adaptation in the negative control (ERKpostStim). To evaluate these phenotypes, we z-scored the features associated to each perturbation to those of the negative control (Figure 4D, see Material and method for details). While many phenotypes were statistically significant, most of them remained mild as observed by visually inspection of the feature distributions (Figure EV3A). Apart from *ERK1+2* KD, only *GRB2, PTK2* and *ERK2* led to a reduction of ERK amplitude (maxPeak). KD of negative regulators such as *SPROUTY 2,3* and *4*, or phosphatases such as *PP2A* and several dual-specificity phosphatases (DUSPs) led to increased ERK amplitude. Increased basal ERK activity was observed for *RKIP, PP2A, DUSP4* and *DUSP6* KDs, indicating a function in regulating basal ERK levels. Prolonged ERK activity (ERKpostStim) was observed in response to KD of *RKIP, PP2A, ERK2, DUSP1,2,3,4,6* and strikingly for *RSK2* KD (Figure EV3B), suggesting a role of these nodes in ERK adaptation.

Because both visual inspection of trajectories, as well as our feature-based approach might miss more subtle ERK dynamics phenotypes, we used CODEX (Jacques et al. 2021), a data-driven approach to identify patterns in single-cell time-series based on convolutional neural networks (CNNs) (Figure EV3C). We trained a CNN to classify ERK trajectories that originate from different siRNA perturbations and selected the ten perturbations for which the CNN classification accuracy was the highest (Appendix Table S4, “CODEX accuracy”, see Material and methods for details). Projection of the CNN features in a t-distributed stochastic neighbor embedding (t-SNE) space revealed different clusters of ERK trajectories (Figure EV3D). Comparison of the ten trajectories with the highest classification confidence identified by CODEX to randomly selected ERK trajectories for low or high optoFGFR expression highlighted ERK phenotypes not accessible to visual inspection and the feature-based approach (Figure 4E). CODEX identified some of the perturbations that affect ERK amplitude, baseline or adaptation observed with the feature-based approach. However, it also highlighted perturbations affecting oscillatory ERK dynamics. *PP2A* KD led to sustained oscillatory behavior. *PLCG1* KD resulted in a first peak followed by damped oscillations, and absence of the dip. As phospholipase C mediates Ca^2+^ signaling in response to FGFR activation (Ornitz and Itoh 2015), this further validates the role of Ca^2+^ signaling in formation of the dip (Appendix Figure S1D-F). *RAPGEF1* KD led to oscillatory ERK responses of different amplitudes. *RSK2, ERK2* and *CRAF* KD displayed reduced oscillatory ERK behavior.

To validate the latter oscillatory ERK dynamics phenotypes, we evaluated the proportion of oscillatory trajectories (trajectories with at least 3 peaks) for each perturbation, both for high and low optoFGFR input (Figure 4F). This confirmed that *RSK2, CRAF* and *ERK2* KD led to decreased oscillatory ERK dynamics. We also observed that these perturbations reduced ERK oscillations in cells stimulated with 1 ng/ml EGF (Figure EV3E-G), suggesting a role of these nodes in the regulation of ERK oscillations in the context of a native RTK.

ERK2 and CRAF isoforms are implicated in the well-established ERK-RAF NFB, known to regulate ERK dynamics (Santos et al. 2007; Ryu et al. 2015; Blum et al. 2019), and to enable consistent ERK dynamics under MEK or ERK perturbations (Fritsche-Guenther et al. 2011; Sturm et al. 2010). *RSK2* encodes the p90 ribosomal S6 kinase 2 protein, an ERK substrate regulating survival and proliferation (Cargnello and Roux 2011; Yoo et al. 2015). RSK2 is also known to be involved in an ERK-induced NFB targeting SOS (Douville and Downward 1997; Saha et al. 2012; Lake et al. 2016), whose significance in the regulation of ERK dynamics has been less well studied. In addition to dampening ERK oscillations, *RSK2* KD also led to slower ERK adaptation when optoFGFR input ceased (Figure 4D, EV3A,B), suggesting an important role of this NFB in ERK dynamics regulation. Our results suggest that the ERK-RAF and ERK-RSK2-SOS NFBs simultaneously operate within the MAPK network to generate ERK oscillations and raise the question whether both NFBs contribute to the strong MAPK signaling robustness observed in our screen.

### Direct optogenetic activation of RAS highlights different ERK dynamics phenotypes than optoFGFR input

To further explore the role of MAPK feedbacks in MAPK signaling robustness, we used optoSOS (Johnson et al. 2017), an optogenetic actuator that activates RAS, and thus bypasses the RSK2-mediated NFB regulation (Figure 5A). OptoSOS consists of a membrane anchored light-activatable iLID domain, and an mCitrine-tagged SspB domain fused to SOS’s catalytic GEF domain. It was stably integrated into cells expressing ERK-KTR and H2B. Because iLID displays faster dissociation rates than CRY2 (t_1/2_= 30 seconds for iLID versus ∼ 5 minutes for CRY2 (Duan et al. 2017; Benedetti et al. 2018)), optoSOS required repeated light pulses to prolong its membrane recruitment and produce a robust ERK response (Figure 5B). Five consecutive 100 ms light pulses at 6 W/cm^2^ (D = 0.6 J/cm^2^) applied at 20-second intervals, provided the minimal light input to induce a saturated ERK amplitude (Figure EV4A). Application of this light input at 2-minute intervals evoked sustained ERK dynamics with small fluctuations at the same frequency as the light input pattern, reflecting the fast optoSOS reversion to the dark state (Figure 5C). OptoSOS did not induce ERK oscillations (Figure EV4B), even in cells with low optoSOS expression or at lower light doses (Figure 5D). However, ERK amplitudes correlated with optoSOS expression level, low optoSOS levels led to low ERK amplitudes, while high actuator expression levels resulted in high ERK amplitudes. Using the minimal light input to trigger saturating ERK amplitude, both optoSOS and optoFGFR led to steep ERK activation and fast adaptation when light stimulation ceased (compare Figures 2C and 5C), as well as similar ERK amplitudes in cells expressing high actuator levels (Figure 5E). However, high optoSOS expression levels moderately increased ERK activity baseline levels in comparison to optoFGFR (Figure EV4C), suggesting that this system is leaky to some extent.

**Figure 5:**
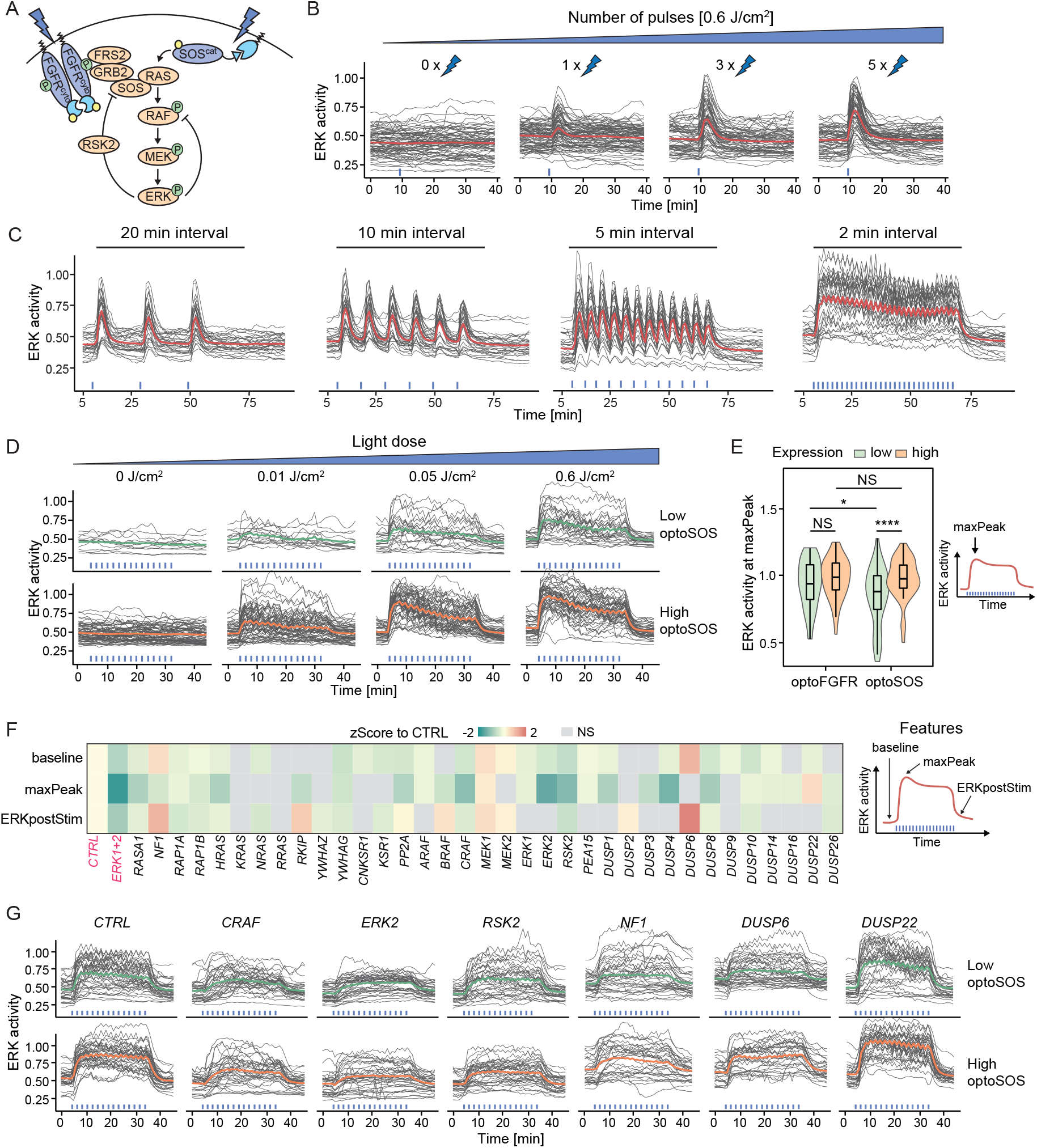
Direct optogenetic activation of RAS highlights different ERK dynamics phenotypes than optoFGFR input. **(A)** Schematic representation of ERK signaling induced by optoSOS versus optoFGFR input. **(B)** ERK dose responses under transient optoSOS input consisting of different numbers of repeated 470 nm pulses (1x, 2x, 3x, 4x and 5x pulses applied at 20-second intervals, D = 0.6 J/cm^2^). Repeated pulses are depicted as a single stimulation (blue bar). **(C)** ERK responses to optoSOS inputs consisting of 5 repeated 470 nm light pulses delivered every 20, 10, 5 and 2 minutes respectively (D = 0.6 J/cm^2^). **(D)** ERK responses to increasing light doses of sustained optoSOS input consisting of 2-minute interval input, each input made of 5 repeated light pulses. Cells were divided in low and high optoSOS expression levels based on the log10 intensity of the optoSOS-mCitrine. **(E)** Quantification of the maxPeak of single-cell ERK responses under sustained optoFGFR (Figure 2F, D = 18 mJ/cm^2^) and optoSOS (Figure 5D, D = 0.6 J/cm^2^) input for low or high expression of each optogenetic system (N = 40 cells per condition). Statistical analysis was done using a Wilcoxon test, comparing each condition to each other (N_min_ = 48 cells per condition, NS: non-significant, *<0.05, **<0.005, ***<0.0005, ****<0.00005, FDR p-value correction method). **(F)** Z-Score evaluation of the baseline, maxPeak and ERKpostStim of single-cell ERK responses under sustained high optoSOS input (D = 0.6 J/cm^2^). The z-score was calculated by comparing each RNAi perturbation to the *CTRL* KD (N_min_ = 33 cells per treatment, from 3 technical replicates). Non-significant (NS) results are in grey (see Figure EV4F for statistical results). **(G)** Single-cell ERK trajectories for low and high optoSOS cells for selected RNAi perturbations (N = 40 randomly selected out of at least 193 trajectories from 3 technical replicates).

Using this specific light input, we performed siRNA screens targeting MAPK signaling nodes downstream of optoSOS in triplicates (Figure EV4D,E). We extracted the baseline, maxPeak, ERKpostStim features from optoSOS high expressing cells (Figure EV4F) and z-scored feature values to the negative control (Figure 5F). We observed more prominent ERK amplitude phenotypes in response to optoSOS input than to optoFGFR input. Some of these phenotypes are shown in Figure 5G. Most prominently, *CRAF, ERK2, DUSP4* KD led to a stronger reduction in ERK amplitude than observed with optoFGFR input. *RSK2* KD also reduced ERK amplitude, suggesting that it also regulates nodes downstream of RAS. However, *RSK2* KD did not decrease ERK adaptation following optoSOS input removal (Figure EV4G), suggesting that it is not involved in NFB regulation in this system. *PP2A* KD did not induce increased ERK amplitude or baseline as observed in the optoFGFR system. As for optoFGFR input, *DUSP6* KD increased basal ERK activity and decreased adaptation (Figure EV4G). *DUSP22* KD led to increased amplitude, without affecting ERK baseline and adaptation. *NF1* KD, which encodes a RAS-specific GAP, led to increased ERK baseline and slower adaptation (Figure EV4G), without affecting ERK amplitude. The NF1 baseline phenotype, that was not observed in the optoFGFR system, might emerge from the optoSOS-mediated low levels of RAS activation due to the optoSOS system’s leakiness (Figure EV4C), that can then be amplified by loss of NF1’s RAS GAP activity. The finding that perturbation of specific nodes (e.g. ERK2 and CRAF) leads to more penetrant phenotypes in response to optoSOS versus optoFGFR input suggested that the RAS/RAF/MEK/ERK part of the network is more sensitive to perturbations than optoFGFR-triggered network, suggesting that the RSK2 NFB that operates above RAS contributes to MAPK signaling robustness.

### Perturbation of the RSK2-mediated NFB increases the efficiency of RAF, MEK and ERK targeting drugs

To further investigate the role of the RSK2-mediated NFB in MAPK signaling robustness, we performed dose response experiments using different MAPK inhibitors and compared ERK amplitudes evoked by optoFGFR (RSK2-feedback dependent) versus optoSOS (RSK2-feedback independent) input, as well as optoFGFR input in absence/presence of RSK2 perturbation. We used drugs targeting B/CRAF (RAF709), MEK (U0126) and ERK (SCH772984). We evaluated the inhibition efficiency by measuring ERK amplitude at a fixed time point, focusing on ERK responses evoked by high optoFGFR or optoSOS inputs to limit the single-cell heterogeneity due to expression variability of the optogenetic actuator. All inhibitors led to a stronger reduction of ERK amplitude and EC_50_ in response to optoSOS versus optoFGFR input (Figure 6A-C, EV5A, Appendix Table S5). Visual evaluation of ERK amplitude distributions (Figure 6B) and quantification of their standard deviations (Figure 6D) revealed more compact ERK amplitude distributions in presence of increasing drug concentrations in response to optoSOS versus optoFGFR input. This suggests a more homogeneous drug inhibition in the cell population in response to optoSOS input. We then performed the identical experiments in *CTRL* or *RSK2* KD cells in response to optoFGFR input (Figure 6E-H, EV5B, Appendix Table S6). *RSK2* KD led to increased inhibition of ERK amplitudes, decreased EC_50_, and more compact ERK amplitude distributions in response to increasing drug concentration than in *CTRL* KD cells. Similar results were observed when the RSK2-mediated feedback was inhibited using the RSK inhibitor SL0101 (Smith et al. 2005) (Figure EV5C-F, Appendix Table S7). Thus, inhibition of the RSK2-mediated NFB sensitizes ERK responses to RAF, MEK or ERK drug perturbations. Note that drug mediated ERK amplitude inhibition was stronger in response to optoSOS input than to optoFGFR input with *RSK2* KD or RSK inhibition, suggesting that additional mechanisms to the RSK2-mediated feedback contribute to MAPK signaling robustness. However, our results suggest that perturbation of the RSK2-mediated feedback can be exploited to enhance the efficiency of MAPK-targeting drugs, reducing ERK amplitudes more homogeneously across the cell population.

**Figure 6:**
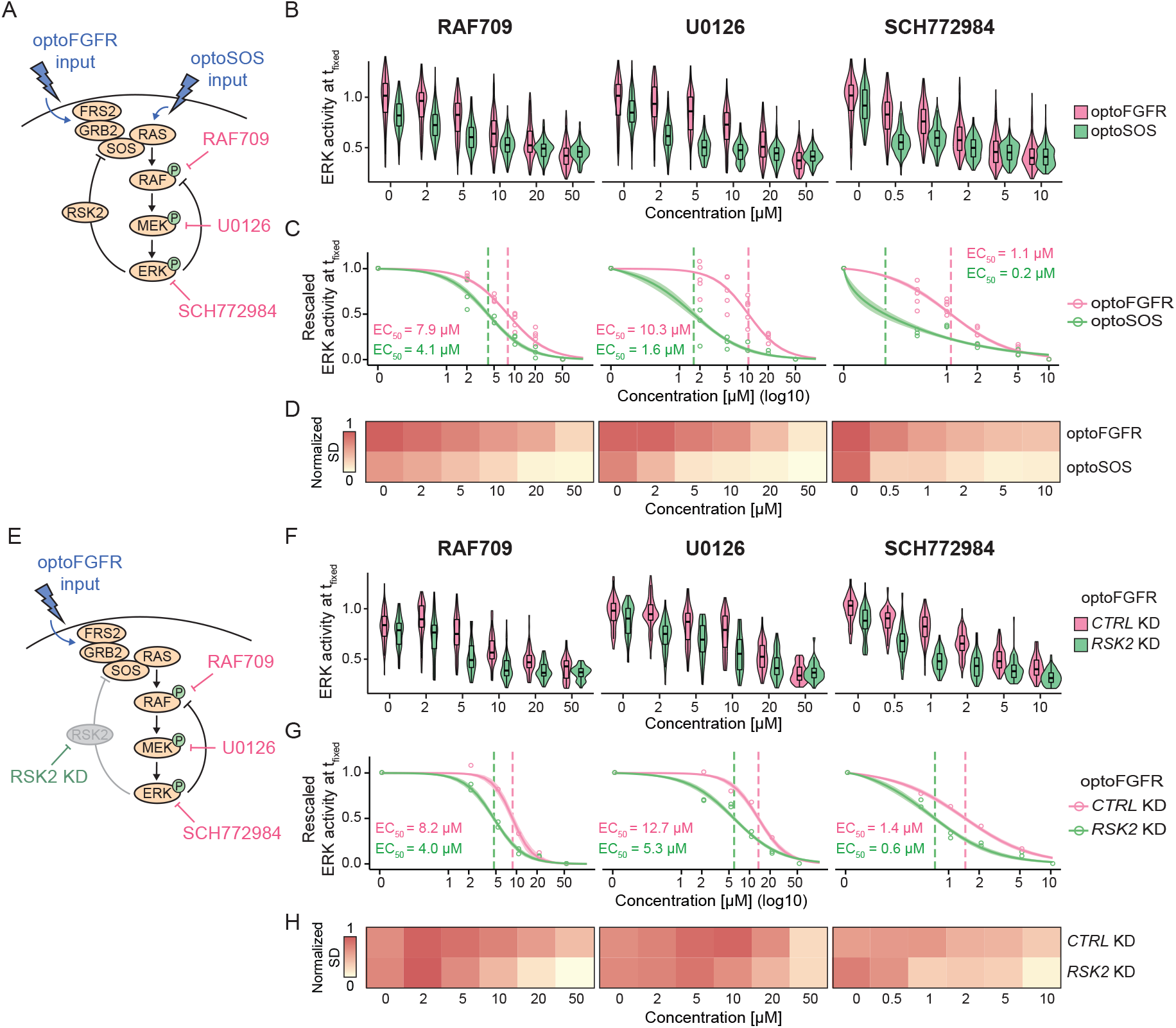
Perturbation of the RSK2-mediated NFB increases the efficiency of RAS, MEK and ERK targeting drugs. **(A)** Schematic representation of the optoFGFR (RSK2-mediated feedback dependent) and optoSOS (RSK2-mediated feedback independent) systems targeted with the B/CRAF (RAF709), the MEK (U0126) or the ERK (SCH772984) inhibitor. **(B)** Single-cell ERK amplitudes from sustained high optoFGFR input (D = 18 mJ/cm^2^) or optoSOS input (D = 0.6 J/cm^2^) under different concentrations of the MAPK inhibitors, extracted at a fixed time point (t_fixed optoFGFR_ = 15 minutes, t_fixed optoSOS_ = 10 minutes, N = 200 cells with high optoFGFR or optoSOS expression per condition randomly selected from 3 technical replicates). **(C)** A Hill function was fit to the normalized mean ERK activity as shown in (B) (N_min_ = 200 cells per condition). Shaded area indicates the 95% CI and dashed lines the EC_50_. **(D)** Normalized standard deviation of ERK amplitudes shown in (B) (N_min_ = 200 cells per condition). **(E)** Schematic representation of the optoFGFR system treated with *CTRL* KD (RSK2-mediated feedback dependent) or *RSK2* KD (RSK2-mediated feedback independent) targeted with the B/CRAF (RAF709), the MEK (U0126) or the ERK (SCH772984) inhibitor. **(F)** Single-cell ERK amplitudes from sustained high optoFGFR input (D = 18 mJ/cm^2^) under different concentrations of the MAPK inhibitors, extracted at a fixed time point (t_fixed optoFGFR_ = 15 minutes, N = 70 cells with high optoFGFR expression per condition (apart from *RSK2* KD + 0 μM U0126 (32 cells), randomly selected from 2 technical replicates for *RSK2* KD and 1 replicate for *CTRL* KD). **(G)** A Hill function was fit to the normalized mean ERK activity as shown in (F) (N_min_ = 32 cells per perturbation). Shaded area indicates the 95% CI and dashed lines the EC_50_. **(H)** Normalized standard deviation of ERK amplitudes shown in (F) (N_min_ = 32 cells per perturbation).

### Targeting the RSK2-mediated feedback in an ErbB2 oncogenic signaling model increases MEK inhibition efficiency

The results above suggested an important role of the RSK2-mediated feedback in MAPK signaling robustness against node perturbation in response to optogenetic inputs in NIH3T3 cells. To test if this feedback also contributes to MAPK signaling robustness in a disease-relevant system, we evaluated its function in MCF10A cells, a breast epithelium model, using either wild-type (WT) or overexpressing ErbB2 (referred to as ErbB2^over^) recapitulating the ErbB2 amplification observed in 20% of all breast cancers (Arteaga and Engelman 2014; Yarden and Pines 2012). We chose this specific model system because ErbB2 amplification leads to constitutive RTK input on the MAPK network, while retaining an intact downstream feedback structure (Figure 7A). This contrasts with other cancer model systems in which additional mutations might lead to RAS or RAF overactivation, and thus disrupt the feedback architecture. Further, previous work has highlighted the role of NFBs in ERK pulse formation in MCF10A cells (Kochańczyk et al. 2017), suggesting that EGFR and ErbB2 trigger a MAPK network with similar feedback circuitry as optoFGFR.

**Figure 7:**
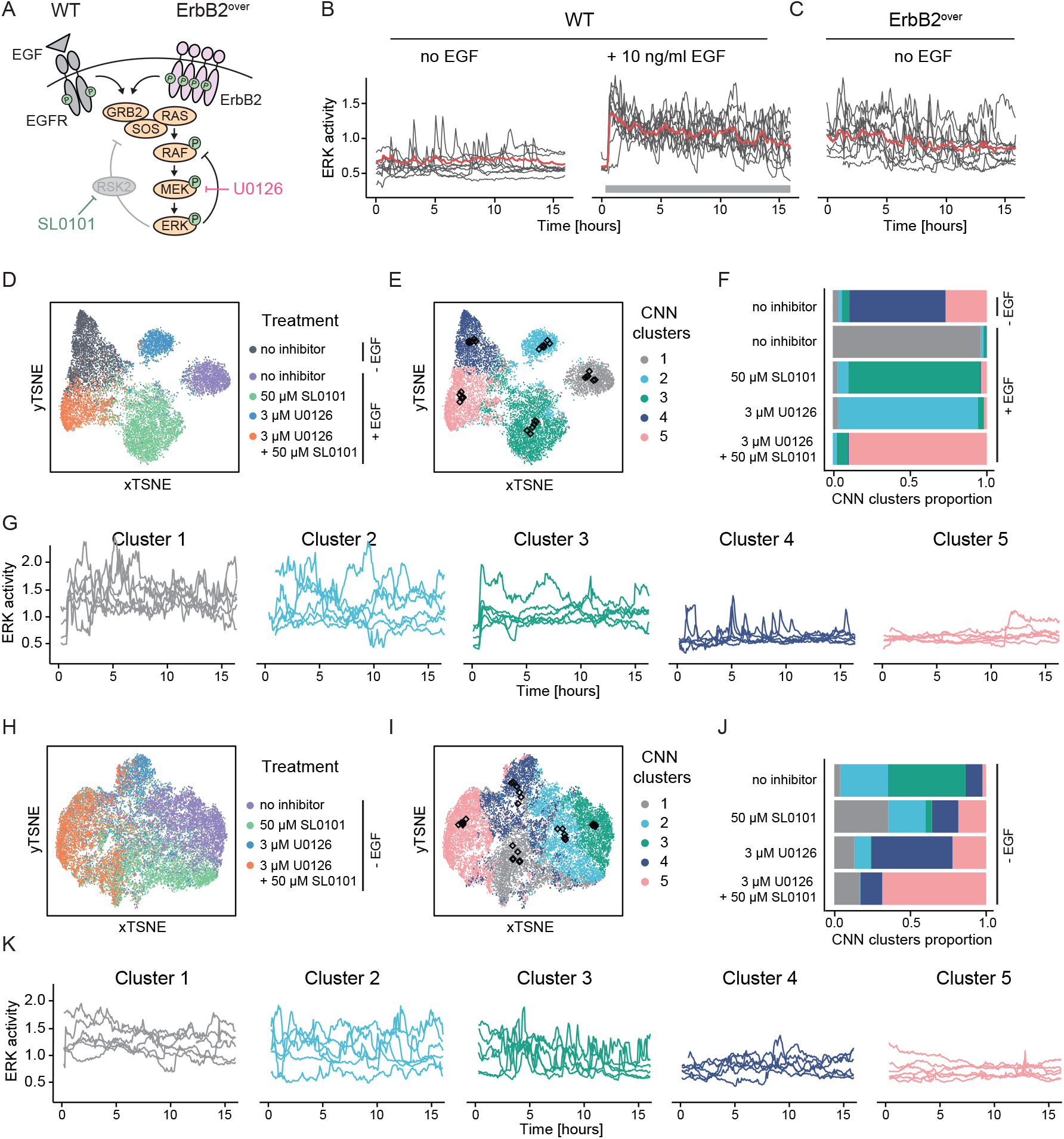
Targeting the RSK2-mediated feedback in an ErbB2 oncogenic signaling model increases MEK inhibition efficiency. **(A)** Schematic representation of MAPK signaling in response to EGFR input in MCF10A WT cells or oncogenic ErbB2 input in ErbB2 overexpressing (ErbB2^over^) cells. **(B-C)** Single-cell ERK responses in MCF10A WT cells without or with stimulation with 10 ng/ml EGF at t = 30 minutes **(B)** and in unstimulated MCF10A ErbB2^over^ cells **(C). (D)** tSNE projection of CODEX’s CNN features from ERK trajectories of MCF10A WT cells without EGF stimulation, or with 10 ng/ml EGF stimulation added at t = 30 min in absence of perturbation, with 50 μM SL0101, 3 μM U0126 or a combination of both. **(E)** t-SNE projection of CODEX’s CNN features shown in (D) colored by the CNN feature clusters. Black diamonds indicate the position of the medoid and its 4 closest neighbor trajectories for each cluster. **(F)** Distribution of the trajectories in the CNN features clusters per treatment. Colors are as shown in (E). **(G)** Medoid trajectories and their 4 closest neighbors per cluster highlighted in (E) (black diamonds). **(H)** tSNE projection of CODEX’s CNN features from ERK trajectories of non-stimulated ErbB2 overexpressing cells without perturbation, with 50 μM SL0101, 3 μM U0126 or a combination of both. **(I)** t-SNE projection of CODEX’s CNN features shown in (H) colored by the CNN feature clusters. Black diamonds indicate the position of the medoid and its 4 closest neighbor trajectories for each cluster. **(J)** Distribution of the trajectories in the CNN features clusters per treatment. Colors are as shown in (I). **(K)** Medoid trajectories and their 4 closest neighbors per cluster highlighted in (I) (black diamonds).

As described before (Albeck et al. 2013), WT cells displayed asynchronous low frequency ERK pulses in the absence of EGF, and high frequency ERK pulses in presence of EGF (Figure 7B). In marked contrast, ErbB2^over^ cells displayed high frequency ERK pulses, even in the absence of EGF (Figure 7C). To investigate the role of the RSK2-mediated feedback in MAPK signaling robustness, we performed a U0126 dose response in EGF-stimulated MCF10A WT cells and found that 3 µM U0126 decreased ERK amplitude without fully suppressing the response (Figure EV5G,H). As observed in response to optogenetic inputs, RSK inhibition with 50 µM SL0101 led to a mild reduction in ERK amplitude. However, in combination with 3 µM U0126, ERK amplitude was decreased to the level of unstimulated cells. Similar results were observed in ErbB2^over^ cells (Figure EV5I), suggesting that RSK2 perturbation increases the sensitivity of ERK responses to MEK inhibition.

As averaging ERK dynamics can hide asynchronous single-cell signaling activity, we further investigated the effect of these perturbations on single-cell trajectories using CODEX (Jacques et al. 2021) (see Material and methods for details). For WT cells, a tSNE projection of the CNN features built from single-cell ERK trajectories hinted that the CNN was able to construct features separating the treatments into well-defined clusters (Figure 7D, EV5J). Clustering of the CNN features confirmed the existence of discrete ERK dynamics clusters (Figure 7E) whose composition correlated with the treatments (Figure 7F). To characterize the dynamics captured by each cluster, we extracted the medoid trajectory and its 4 closest neighbors from each cluster (Figure 7G). This revealed that non-stimulated cells mostly display low frequency ERK activity pulses (cluster 4) or absence of pulses (cluster 5). Cells stimulated with EGF without inhibitor displayed ERK pulses of high amplitude (cluster 1). SL0101-treated cells displayed a sustained ERK activation at low amplitude (cluster 3). U0126-treated cells still displayed prominent ERK pulses but at a lower amplitude than EGF-treated cells in absence of drug (cluster 2). Finally, in cells treated with both U0126 and SL0101, almost no ERK activity was observed (cluster 5). For ErbB2^over^ cells, we observed that the CNN features were forming a more continuous space with less distinct clusters (Figure 7H,I, EV5K). A heterogeneous mix of ERK trajectory clusters was observed for the different treatments (Figure 7J,K). Untreated cells mostly displayed high frequency ERK pulses that were either sharp (cluster 3) or wider (cluster 2). SL0101-treated cells were almost equally shared between cluster 1 (relatively flat high amplitude ERK trajectories), cluster 2, cluster 4 (low amplitude ERK pulses) and cluster 5 (low baseline activity). U0126 led to a less heterogeneous mix mostly consisting of ERK trajectories from cluster 4 and 5. The U0126/SL0101 combination shifted most cells to cluster 5, indicating an efficient inhibition of ERK activity at a suboptimal U0126 concentration. Thus, RSK inhibition also sensitizes the MAPK network to U0126-mediated MEK inhibition both in MCF10A WT and ErbB2^over^ cells.

## Discussion

### Optogenetic actuator-biosensor circuits allow for feedback structure mapping in the MAPK network

ERK dynamics is crucial for fate decisions. Yet, the topology of the network enabling the cells to sense different inputs and convert this information into finely tuned ERK dynamics remains poorly understood. We developed genetic circuits consisting of optogenetic actuators and an ERK biosensor (Figure 1A, 5A) that allow for a large-scale interrogation of single-cell ERK dynamics and investigated the effects of 50 RNAi perturbations targeting components of the MAPK signaling network (Figure 4A). In our optoFGFR screen, we only observed a small number of penetrant ERK dynamics phenotypes (Figure 4D-F), implying that the MAPK network can buffer against perturbations of most of its components. We cannot exclude that in some cases, even on the relatively short 72 hours timescale of the RNAi experiment, compensation by upregulation of specific nodes might occur. However, our data suggest that the MAPK network topology allows for MAPK signaling robustness – the production of consistent ERK outputs in presence of node perturbation. This might emerge from isoform redundancy for multiple nodes in the network, as observed for single or combined ERK isoforms perturbation (Figure 4B), but also for individual perturbation of RAS, RAF, MEK isoforms. Another mechanism might involve NFBs that have been shown to decrease the network sensitivity to node perturbation (Sturm et al. 2010; Fritsche-Guenther et al. 2011). Our screen suggested that RSK2, that mediates a NFB from ERK to SOS (Douville and Downward 1997; Saha et al. 2012), both regulates ERK dynamics (Figure 4D-F) and plays a role in MAPK signaling robustness (Figure 6E-H). In addition, our data suggest that the well-studied ERK-RAF NFB, which has been shown to buffer against MAPK node perturbations (Sturm et al. 2010; Fritsche-Guenther et al. 2011), also regulates ERK dynamics (Figure 4F). We therefore speculate that both feedbacks operate simultaneously in the MAPK network, and act at multiple levels within the cascade to warrant MAPK signaling robustness. Consistently with this hypothesis, we observed that the optoSOS-triggered network, which is not under the RSK2 NFB regulation, shows an increased sensitivity in ERK amplitude to perturbation of some nodes (Figure 5F,G). Indeed, ERK2 and CRAF perturbations, which led to loss of ERK oscillations, had relatively mild amplitude phenotypes in response to optoFGFR input, while both perturbations led to strong ERK amplitude phenotypes in response to optoSOS input. Because these phenotypes were not observed with other ERK and RAF isoforms, we propose that ERK2 and CRAF are the isoforms involved in the classic ERK-RAF NFB. Additional feedbacks have been reported within the MAPK network (Langlois et al. 1995; Lake et al. 2016; Kochańczyk et al. 2017), and even if they have not been highlighted in our screen, they might also regulate ERK dynamics.

While providing the experimental throughput to perturb and analyze ERK dynamics at scale, optoFGFR, that lacks an ectodomain, evoked different ERK dynamics than endogenous RTKs such as FGFR and EGFR (Figure 3A,B compared to Figure 2F). These different ERK dynamics emerge likely because of receptor-level interactions that involve competition of bFGF for FGFR and heparan sulfate proteoglycan co-receptors (Kanodia et al. 2014; Blum et al. 2019) in the case of FGFR, or receptor endocytosis in the case of EGFR (Kiyatkin et al. 2020; Gerosa et al. 2020). Our combined modeling and experimental approach suggested that optoFGFR and EGFR share similar downstream MAPK network circuitries and NFBs (Figure 3C-G). OptoFGFR therefore provides a simplified system that allowed us to focus on intracellular feedback structures, without confounding receptor level regulations. Our Bayesian inference modeling approach, that is parameter agnostic, could provide simple intuitions about the receptor-level and negative feedback structures that shape ERK dynamics in response to optoFGFR and EGFR inputs. However, even if we had access to many ERK dynamics phenotypes, our modeling approach did not allow us to explore more sophisticated MAPK network topologies such as the presence of two NFBs or multiple node isoforms. We interpreted our data using some of the feedback structures that have been previously experimentally documented and modelled but cannot formally exclude that the observed ERK dynamics emerge from different network structures. In the future, information about the different nodes and their dynamics might allow to further constrain the model topology and parameter space, and hopefully address this limitation.

### Additional novel insights into regulation of ERK dynamics

Our optoFGFR and optoSOS screens provided new system-wide insights into the regulation of the MAPK network. Strikingly, the same perturbations induced different ERK dynamics phenotypes in the optoFGFR and optoSOS screens. This might occur because some regulators target the MAPK network at multiple levels, differently affecting ERK responses triggered with optoFGFR or optoSOS inputs. Additionally, as the two optogenetic systems are under the regulation of one versus two simultaneously occurring NFBs, they might have different sensitivities to perturbations, as discussed above.

With respect to the optoFGFR system, *GRB2* KD led to a reduction of ERK amplitude (Figure 4D,E). GRB2 acts as the RTK-proximal adaptor to activate SOS (Chardin et al. 1993; Belov and Mohammadi 2012). As GRB2 operates at the start of the cascade, outside of most NFBs, heterogeneity in its expression levels might be less easily buffered out. *PLCG1* KD increased damped oscillatory behavior (Figure 4E,F). Phospholipase Cɣ1 activates calcium signaling, which has itself been shown to regulate RAS/MAPK signaling in a calcium spike frequency-dependent manner (Kupzig et al. 2005; Cullen and Lockyer 2002). Further investigation will be required to understand the significance of this crosstalk. *RKIP* KD resulted in higher ERK baseline and slower ERK adaptation post stimulation, without affecting ERK amplitude (Figure 4D). RKIP (RAF kinase inhibitor protein) prevents MEK phosphorylation by CRAF (Yeung et al. 2000), suggesting that RKIP-dependent regulation is specifically involved in keeping basal ERK activity low. With respect to phosphatases, none of their perturbations led to a strong phenotype such as sustained ERK dynamics post stimulation for example. The strongest phenotype was observed for *PP2A* KD that led to increased ERK amplitude, baseline, and slower adaptation (Figure 4D, EV3A). This might occur because the protein phosphatase 2A is an ubiquitous phosphatase that acts at multiple levels by dephosphorylating SHC1, MEK1, MEK2, ERK1 and ERK2, as well as a large number of other proteins (Junttila et al. 2008; Saraf et al. 2010). The observation that in optoFGFR-low *PP2A* KD cells, ERK dynamics displayed increased amplitude but still oscillated rather than exhibiting sustained behavior, suggests that NFBs might buffer against the loss of phosphatase regulation to some extent. Perturbation of the nuclear DUSPs, DUSP1,2,4, the atypical DUSP3 and most strongly the cytosolic DUSP6 (Patterson et al. 2009) led to higher ERK baseline, reduced adaptation, with only limited effects on amplitude (Figure 4D, EV3A). Consistently, DUSP6 has previously been proposed to pre-emptively dephosphorylate MAPKs to maintain low ERK activity baseline levels at resting state (Huang and Tan 2012). Our results indicate that perturbation of single DUSPs might not be compensated by the others, suggesting that individual DUSPs might regulate specific substrates within the MAPK network. Except for *DUSP6*, KD of the different DUSPs did not significantly affect oscillatory ERK behavior in optoFGFR-low cells (Figure 4F), suggesting that they are not involved in the MAPK feedback circuitry that operates on timescales of minutes.

The optoSOS screen revealed stronger ERK amplitude phenotypes, especially for *ERK2* and *CRAF* KD (Figure 5F versus 4D). Unlike for optoFGFR input, *RSK2* KD did not result in slower ERK adaptation, suggesting that ERK responses triggered by the optoSOS input are not regulated by the RSK2-mediated NFB. However, *RSK2* KD led to a reduction of ERK amplitude, also observed to a lesser extent in response to optoFGFR input, suggesting a role of RSK2 in ERK amplitude regulation downstream of RAS. With respect to phosphatases, *PP2A* KD led to decreased amplitude, a different phenotype than in response to optoFGFR input. This might occur because of the broad specificity PP2A phosphatase, which might lead to different phospho-proteomes in response to optoSOS versus optoFGFR input. Similar phenomena might apply for most of the DUSPs.

### The RSK2-mediated feedback can be targeted to potently inhibit oncogenic ErbB2 signaling

Our data suggest that the RSK2-mediated NFB is important for MAPK signaling robustness downstream of our prototypic optoFGFR RTK (Figure 6). We found that the RSK2-mediated NFB likely also operates downstream of EGFR and oncogenic ErbB2 signaling in MCF10A cells (Figure 7). In response to EGF stimulation, or ErbB2 overexpression, a subset of RSK-inhibited cells displayed wider ERK pulses, suggesting that the RSK2 NFB is also involved in ERK adaptation in this system (Figure 7G cluster 3, Figure 7K cluster 1 and 2). Further, RSK inhibition led to a high heterogeneity of ERK dynamics within the cell population especially visible in the case of ErbB2 overexpressing cells (Figure 7J), which might result from the reduced ability of the MAPK network to cope with nodes expression noise in absence of the RSK2 NFB. In EGF-treated cells, combination of RSK and suboptimal MEK inhibition led to strong and homogeneous ERK inhibition (Figure 7E-G, cluster 5). In the ErbB2 overexpressing cells, combined RSK/MEK inhibition shifted most of the cell population to flat, low amplitude ERK dynamics, enabling to further inhibit a large number of cells when compared to suboptimal MEK inhibition only (Figure 7I-K, cluster 5). These results suggest that pharmacological inhibition of the RSK2-mediated NFB can be used to reduce MAPK signaling robustness, sensitizing the network to MEK perturbation. Such non-trivial drug combinations might allow for homogeneous inhibition of ERK dynamics in most of the cells in a population. This homogeneous inhibition might mitigate the emergence of drug-tolerant persister cells from cell subpopulations that display residual ERK activity in response to inhibition of a single node. Our results imply that efficient pharmacological inhibition of the MAPK network requires precise understanding of its topology. The RSK2 NFB is an example of a druggable node that can be exploited to target MAPK signaling robustness.

Our scalable experimental pipeline provides new insight into the MAPK network wiring that produces ERK dynamics. However, our perturbation approach only highlighted very subtle ERK dynamics phenotypes, precluding a complete understanding of the MAPK network. We envision that this will require more precise knowledge about the dynamics of MAPK network nodes and their interactions in response to defined inputs and perturbations. Such data can now be produced at scale using optogenetic actuator/biosensor circuits as those we describe in this work. This information might allow for faithful parametrization of more complex models. With the increasing amount of optogenetic actuators and biosensors available, similar genetic circuits could also be designed to study the dynamics of other signaling pathways at scale.

## Materials and methods

### Cell culture and reagents

NIH3T3 cells were cultured in DMEM high glucose medium with 5% fetal bovine serum, 4 mM L-glutamine, 200 U/ml penicillin and 200*µ*g/ml streptomycin at 37°C with 5% CO_2_. All imaging experiments with NIH3T3 were done in starving medium consisting of DMEM high glucose supplemented with 0.5% BSA (Sigma), 200 U/ml penicillin, 200 μg/ml streptomycin and 4 mM L-Glutamine. MCF10A human mammary cells were cultured in DMEM:F12 supplemented with 5% horse serum, 20 ng/ml recombinant human EGF (Peprotech), 10 μg/ml insulin (Sigma), 0.5 μg/ml hydrocortisone (Sigma), 200 U/ml penicillin and 200 μg/ml streptomycin. All imaging experiments with MCF10A were done in starving medium consisting in DMEM:F12 supplemented with 0.3% BSA, 0.5 μg/ml hydrocortisone, 200 U/ml penicillin and 200 μg/ml streptomycin. For growth factor stimulations, we used human EGF (AF-100, Peprotech) and human basic FGF (F0291, Sigma). Chemical perturbations were done with SU-5402 (SML0443, Sigma), RAF709 (HY-100510, Lucerna Chem), U0126 (S1102, Selleck chemicals, Lubio), SCH772984 (HY-50846, Lucerna-Chem), SL0101 (559285, Sigma), Cyclosporine A (10-1119, Lucerna-chem) and Ionomycin (sc-3592, Santa Cruz). Selection of the cells post transfection was done using Puromycin (P7255, Sigma), Blasticidin S HCI (5502, Tocris) and Hygromycin B (sc-29067, Lab Force).

### Plasmids and stable cell line generation

The optoFGFR construct was a gift from Won Do Heo (Addgene plasmid # 59776) (Kim et al. 2014). It consists of the myristoylated FGFR1 cytoplasmic region fused with the PHR domain of the cryptochrome2 and tagged with mCitrine. It was cloned in a lentiviral backbone for stable cell line generation. A modified version of the optoFGFR tagged with the red fluorophore mScarlet (Bindels et al. 2017) was cloned in a PiggyBac plasmid pPBbSr2-MCS (blasticidin resistance), a gift from Kazuhiro Aoki. The optoSOS construct is a modified version of the tRFP-SSPB-SOScat-P2A-iLID-CAAX (Addgene plasmid #86439) (Johnson et al. 2017), in which we replaced the tRFP by mCitrine. The construct was cloned in the pPB3.0.Puro, an improved PiggyBac plasmid generated in our lab with puromycin resistance. The ERK-KTR-mRuby2 and ERK-KTR-mTurquoise2 reporters were generated by fusing the ERK Kinase Translocation Reporter (ERK-KTR) (Regot et al. 2014) with mRuby2 (Lam et al. 2012) or mTurquoise2 (Goedhart et al. 2012). The nuclear marker H2B-miRFP703 is a fusion of the human H2B clustered histone 11 (H2BC11) with the monomeric near-infrared fluorescent protein miRFP703 (Shcherbakova et al. 2016) (Addgene plasmid #80001). ERK-KTR-mRuby2, ERK-KTR-mTurquoise2 and H2B-miRFP703 were cloned in the PiggyBac plasmids pPB3.0.Hygro, pSB-HPB (gift of David Hacker, EPFL, (Balasubramanian et al. 2016)) and pPB3.0.Blast, respectively. All constructs in PiggyBac plasmids were co-transfected with the helper plasmid expressing the transposase (Yusa et al. 2011) for stable insertion using the jetPEI (Polyplus) transfection reagent for NIH3T3 cells or FuGene (Promega) transfection reagent for MCF10A cells. After antibiotic selection, NIH3T3 cells were FACS-sorted to generate stable cell lines homogeneously expressing the biosensors. In the case of MCF10A cells, clones with uniform biosensor expression were isolated. To generate ErbB2 overexpressing MCF10A cells, lentiviral transduction using a pHAGE-ERBB2 construct (a gift from Gordon Mills & Kenneth Scott, Addgene plasmid #116734, (Ng et al. 2018)) was performed in the presence of 8 μg/ml polybrene (TR1003, Sigma) in cells already expressing H2B-miRFP703 and ERK-KTR-mTurquoise2. Cells were further selected with 5 μg/ml puromycin.

### Live imaging of ERK dynamics

NIH3T3 cells were seeded in 96 well 1.5 glass bottom plates (Cellvis) coated with 10 μg/ml Fibronectin (Huber lab) using 1.5 × 10^3^ cells/well and incubated for 24 hours. MCF10A cells were seeded in 24-well 1.5 glass bottom plates (Cellvis) coated with 5 μg/ml Fibronectin (Huber lab) at 1 × 10^5^ cells/well and incubated for 48 hours. NIH3T3 cells were washed with PBS and incubated in starving medium for 4 hours in the dark before starting the experiment. MCF10A cells were starved for 7 hours before starting the experiments. In experiments involving drug perturbations, cells were incubated for 2 hours (or 1 hour in MCF10A experiments) with the inhibitor(s). Imaging was performed with an epifluorescence Eclipse Ti inverted fluorescence microscope (Nikon) using a Plan Apo air 20x (NA 0.8) objective. Nikon Perfect Focus System (PFS) was used to keep cells in focus throughout the experiment. Illumination was done with a SPECTRA X light engine (Lumencor) with the following filters (Chroma): mTurquoise2: 440 nm LED, 470lp, 69308 CFP/YFP/mCherry-ET, CFP 458-482; mCitrine: 508 nm LED, ET500/20x, 69308bs, ET535/30m; mRuby2 and mCherry: 555 nm LED, ET575/25x, 69008bs, 59022m, miRFP703: 640 nm LED, ET640/30x, 89100bs Sedat Quad, 84101m Quad. Images were acquired with an Andor Zyla 4.2 plus camera at a 16-bit depth. Image acquisition and optogenetic stimulation were controlled with the NIS-Element JOBS module. For NIH3T3 experiments, ERK-KTR-mRuby2 and H2B-miRFP703 were acquired at 1-minute interval and 470 nm light inputs were delivered at specific frequencies and intensities (see below). MCF10A image acquisition was performed at 5-minute time resolution. Growth factor stimulations were done by manually pipetting EGF and bFGF during the experiment. We used mCitrine intensity to quantify the expression level of the optogenetic constructs. However, as mCitrine excitation leads to optoFGFR or optoSOS activation, we acquired one frame with the ERK-KTR-mRuby2, the H2B-miRFP703 and the mCitrine-tagged optoFGFR or optoSOS only at the end of each NIH3T3 experiments. All experiments were carried on at 37°C with 5% CO_2_.

### Optogenetic stimulation

Light stimulations were delivered with a 470 nm LED light source that was hardware-triggered by the camera to generate light pulses of reproducible duration. Light stimulations of defined intensity and duration were programmed to be automatically delivered at specific timepoints. To define the dose of light received by the cells, we measured the 470 nm light intensity at the focal plane using an optical power meter (X-Cite Power Meter, Lumen Dynamics Group) and converted this value to a power density as

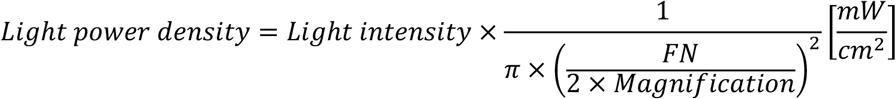

with FN = 18 mm. The obtained value was then multiplied by the duration of the pulse to obtain the dose of light received by the cells for each light pulse.

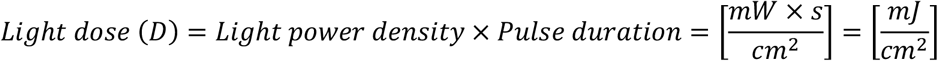

For stimulation of the optoFGFR cells, the 470 nm LED intensity was limited to a low dose by combining a ZET470/10x filter and a ND filter 5% (Chroma). Transient stimulations were done with a single pulse, while sustained stimulations were done with single pulses delivered every 2 minutes. For stimulation of the optoSOS cells, we used the 470 nm LED with a ET470/24x filter (no ND filter). Transient stimulations were done with 5 pulses repeated at 20-second intervals, while sustained stimulations were done using 5 pulses repeated at 20-second intervals, delivered every 2 minutes.

| Figures | System | Power density | Pulse duration | Dose | Stimulation pattern |
| --- | --- | --- | --- | --- | --- |
| 1B,C, Appendix S1A, Appendix S1D | optoFGFR | 180 mW/cm<sup>2</sup> | 1 x 100 ms | 18 mJ/cm<sup>2</sup> | transient |
| Appendix S1B,C | optoFGFR (mScarlet) | > 180 mW/cm<sup>2</sup> | 1 x 100 ms | > 18 mJ/cm<sup>2</sup> | transient |
| 2A,B | optoFGFR | variable | variable | variable | transient |
| 2C, EV1A,B | optoFGFR | 180 mW/cm<sup>2</sup> | 1 x 100 ms | 18 mJ/cm<sup>2</sup> | variable |
| 2D,E, 3C,E, EV1K, 4B,D-F, EV2C,D, EV3A-D, 5E, EV4C, 6B-D,F-H, EV5A,B,D-F | optoFGFR | 180 mW/cm<sup>2</sup> | 1 x 100 ms | 18 mJ/cm<sup>2</sup> | sustained |
| 2F | optoFGFR | variable | 1 x 100 ms | variable | sustained |
| 5B | optoSOS | 6 W/cm<sup>2</sup> | variable x 100 ms (20 sec interval) | 0.6 J/cm<sup>2</sup> | transient |
| EV4A | optoSOS | variable | variable x 100ms (20-sec interval) | variable | transient |
| 5C | optoSOS | 6 W/cm<sup>2</sup> | 5 x 100 ms (20- | 0.6 J/cm<sup>2</sup> | variable |
|  |  |  | sec interval) |  |  |
| 5D | optoSOS | variable | 5 x 100 ms (20-sec interval) | variable | sustained |
| 5E-G, EV4B-G, 6B-D, EV5A | optoSOS | 6 W/cm<sup>2</sup> | 5 x 100 ms (20-sec interval) | 0.6 J/cm<sup>2</sup> | sustained |

### TIRF imaging of optoFGFR dynamics

Cells were seeded at a density of 1 × 10^3^ per well in 96 well 1.5 glass bottom plates (Cellvis) coated with 10 μg/ml Fibronectin (Huber lab) and incubated for 24 hours at 37°C with 5% CO_2_. Before imaging, cells were washed with PBS and incubated in starving medium for 4 hours in the dark. Imaging was performed with an epifluorescence Eclipse Ti inverted fluorescence microscope (Nikon) using a CFI Apochromat TIRF 100x oil (NA 1.49). Images were acquired with an Andor Zyla 4.2 plus camera at a 16-bit depth. TIRF images were acquired with a 561 nm laser using a ET575/25 filter in front of the ZT488/561rpc (Chroma) to prevent nonspecific activation of the CRY2. MetaMorph software (Universal Imaging) was used for acquisition. TIRF images of the optoFGFR-mScarlet were acquired at a 20-second interval. Optogenetic stimulation was done using a 470 nm LED (SPECTRA X, Lumencor) (Appendix Figure S1B). All experiments were carried on at 37°C with 5% CO_2_.

### Image processing pipeline

Nuclear segmentation was done in CellProfiler 3.0 (McQuin et al. 2018) using a threshold-based approach of the H2B channel. In the case of MCF10A cells, nuclear segmentation was preceded by prediction of nuclear probability using a random forest classifier based on different pixel features available in Ilastik software (Berg et al. 2019). To measure the ERK-KTR fluorescence in the cytosol, the nuclear mask was first expanded by 2 pixels to exclude the blurred edges of the nucleus. The new mask was then further expanded by 4 pixels in a threshold-based manner to obtain a “ring” area corresponding to the cytoplasmic ERK-KTR. ERK activity was obtained by calculating the ratio between the average cytosolic pixel intensity and the average nuclear pixel intensity. Single-cell tracking was done on nuclear centroids with MATLAB using μ-track 2.2.1 (Jaqaman et al. 2008). The final images containing the ERK-KTR-mRuby2, H2B-miRFP703 and the optoFGFR-mCitrine (or optoSOS-mCitrine) channels were processed using the same CellProfiler settings as the time lapse images. Intensity of the mCitrine was extracted under the ERK-KTR cytoplasmic mask and used to classify cells into low or high expressors in a threshold-based manner. For optoFGFR-evoked ERK responses, the threshold was defined empirically to separate oscillatory and non-oscillatory ERK responses (low < -1.75 (log10 mCitrine intensity) < high). For optoSOS-evoked ERK responses, the threshold was defined empirically to separate cells with low or high ERK response amplitudes (low < -1.25 (log10 mCitrine intensity) < high). The same thresholds were kept across experiments to compare low and high expressors.

The optoFGFR-mScarlet dimers/oligomers were segmented using the pixel classification module from Ilastik (Berg et al. 2019). OptoFGFR dimers, cell background and trafficking vesicles were manually annotated on images before and after the light stimulation. A probability map of the optoFGFR dimers classification was exported as TIFF for each frame. We then computed the mean of pixel intensities from the binarized mask obtained with Ilastik using Fiji (Appendix Figure S1C).

### Quantification of ERK activity

We wrote a set of custom R scripts to automatically calculate the ERK-KTR cytoplasmic/nuclear ratio as a proxy for ERK activity for each single-cell, link single-cell ERK responses with the corresponding optoFGFR/optoSOS intensity value and export the corresponding ERK single-cell trajectories. For NIH3T3 data, outliers in ERK single-cell trajectories were removed using a clustering-based approach (https://github.com/pertzlab/Outlier_app). Trajectories with an ERK activity higher than 0.8 or lower than 0.2 before stimulation, above 1.6 during the whole experiment or displaying single time point spiking values were removed. For MCF10A data, trajectories with an ERK activity above 2 or shorter than 90% of the total experiment duration were removed. All the R codes used for further analysis are available as supplementary information (see Data availability section). Hierarchical clustering analysis of single-cell trajectories (Figure 2D, EV3F,G, EV4B) was done using Time Course Inspector (Dobrzyński et al. 2019).

### Modeling

The model for the EGF and light stimulated ERK cascade is a kinetic model, representing the EGF receptor, the inter-cellular proteins (RAS, RAF, MEK, ERK) as well as a negative feedback (NFB) from ERK to RAF and the inactivation of the EGF receptor in the form of endocytosis (Figure 3D). We explicitly modelled the ERK-KTR readout through nuclear and cytosolic KTR. The initial fraction of cytosolic KTR is estimated from the data through the parameter *ktr*_*init*_. The KTR readout *Y*(*t*) was taken to be the ratio of cytosolic KTR over nuclear KTR with additive Gaussian noise

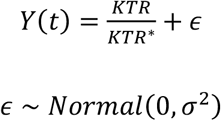

where the variance of the measurement noise *σ*^2^ was estimated from the data. Appendix Table S1 shows all modelled species, their notation used for the equation, as well as the initial values. We assume that in the beginning of the experiment, all species are in the inactive form, reflecting the fact that the cells have been starved. The total concentrations of all species have been normalized to 1. The model equations are shown in Appendix Table S2. The phosphorylation events are modeled with Michaelis-Menten kinetics. The NFB is modelled through the modeling species *NFB* and its “active” version *NFB ∗* which affects the dephosphorylation rate of *RAF* linearly. The activation, endocytosis, and recycling of the EGF receptor is modelled linearly. The model parameters are described in Appendix Table S3. For the modeling of the two smaller models (without feedback (Figure EV1E) or without endocytosis (Figure EV1H)), we set the corresponding parameters (*k*_*nfb*_ and *r*_2,3_) to zero.

For the parameter inference, we used a Nested Sampling algorithm as described in (Mikelson and Khammash 2020). The inference was performed on the ETH High-performance Cluster Euler and was done using the parallel implementation on 48 cores. The algorithm was run for 24 hours or until the algorithm stopped because the termination criterion *Δ*_*LFNS*_ (see (Mikelson and Khammash 2020) for details) was −∞. As prior distributions, we chose for all parameters non-informative log-uniform priors between 10^−5^ and 10^5^, except for *ktr*_*init*_ for which we chose a uniform prior on the interval [0, 1] and for σ for which we chose a log-uniform prior between 10^−5^ and 1. Predictive distributions can be found on Figure 3F,G, EV1F,G,I,J.

### RNAi perturbation screen

We used Ingenuity Pathway Analysis (IPA, Qiagen) to select proteins directly interacting with ERK, MEK, RAF, RAS and FGFR, that are known to be expressed in NIH3T3 cells using a proteomics approach (Schwanhäusser et al. 2011; Jensen et al. 2009) (Appendix Table S4). We then imported this protein list in STRING (Jensen et al. 2009) to generate an interaction network with a minimum interaction score of 0.4. The final interactome was manually modified to display the protein names to facilitate the readout (Figure 4A). We targeted these selected proteins with RNA interference, using the siPOOL technology (one siPOOL containing a mix of 30 siRNAs targeting the same gene (Hannus et al. 2014), sequences available in the Data availability section). We arranged the siPOOLs in a 96 well plate format (in columns 2-5 and 8-11, one well per siPOOL) with the non-targeting siRNA (*CTRL*) and the positive control (mix of 5 nM siPOOL against *ERK1* and 5 nM siPOOL against *ERK2*) placed alternately in columns 1, 6, 7 and 12. Cells were reverse transfected using RNAiMAX (Thermofisher, 13778150) following the recommended siPOOL transfection protocol (https://sitoolsbiotech.com/protocols.php). OptoFGFR-expressing cells were transfected with 10 nM of siPOOL in a 96 well 1.5 glass bottom plate (Cellvis) coated with 10 μg/ml Fibronectin (Huber Lab) at 0.3 × 10^3^ cells/well density and incubated for 72 hours at 37°C and 5% CO_2_. For the imaging, the 96 well plate was divided into 15 sub-experiments, each sub-experiment consisting of a negative control well, a positive control well and 4 wells with different siPOOLs. We selected 2 FOVs per well and programmed the microscope to run the 15 experiments sequentially, acquiring the ERK-KTR-mRuby2 and the H2B-miRFP703 channels with a 1-minute interval, stimulating the cells with sustained optoFGFR input (2-minute intervals, D = 18 mJ/cm^2^), and acquiring a final frame with ERK-KTR-mRuby2, H2B-miRFP703 and optoFGFR-mCitrine (Figure 4B,D-F, EV2C,D, EV3A-D, 6F-H, EV5B). For the optoSOS system, we limited the perturbation screen to targets acting below RAS (Figure 5F,G, EV4D-G). Stimulations were done with sustained optoSOS input (5 repeated pulses at 2-minute intervals, D = 0.6 J/cm^2^). For EGF experiments, cells were stimulated with 1 ng/ml EGF at t = 5 minutes (Figure EV3E-G).

### Real-time qPCR

Cells were transfected with different concentrations of siPOOL in a 24 well plate at 5 × 10^3^ cells/well density and incubated at 37°C with 5% CO_2_ for 72 hours before RNA isolation. Reverse transcription was done with the ProtoScript II reverse transcriptase kit (Bioconcept, M0368L). Real-time qPCR reactions were run using the MESA Green pPCR MasterMix Plus for SYBR Green assay (Eurogenetec, RT-SY2X-03+WOU) on the Rotor-Gen Q device (Qiagen). Each sample was tested in triplicate. Expression level of the gene of interest was calculated using the 2^-ΔΔCt^ method with *GAPDH* expression level as internal control (Figure EV2A). The following primers were used for the RT-qPCR reaction (designed with the Real-time PCR (TaqMan) Primer and Probes Design Tool from GenScript).

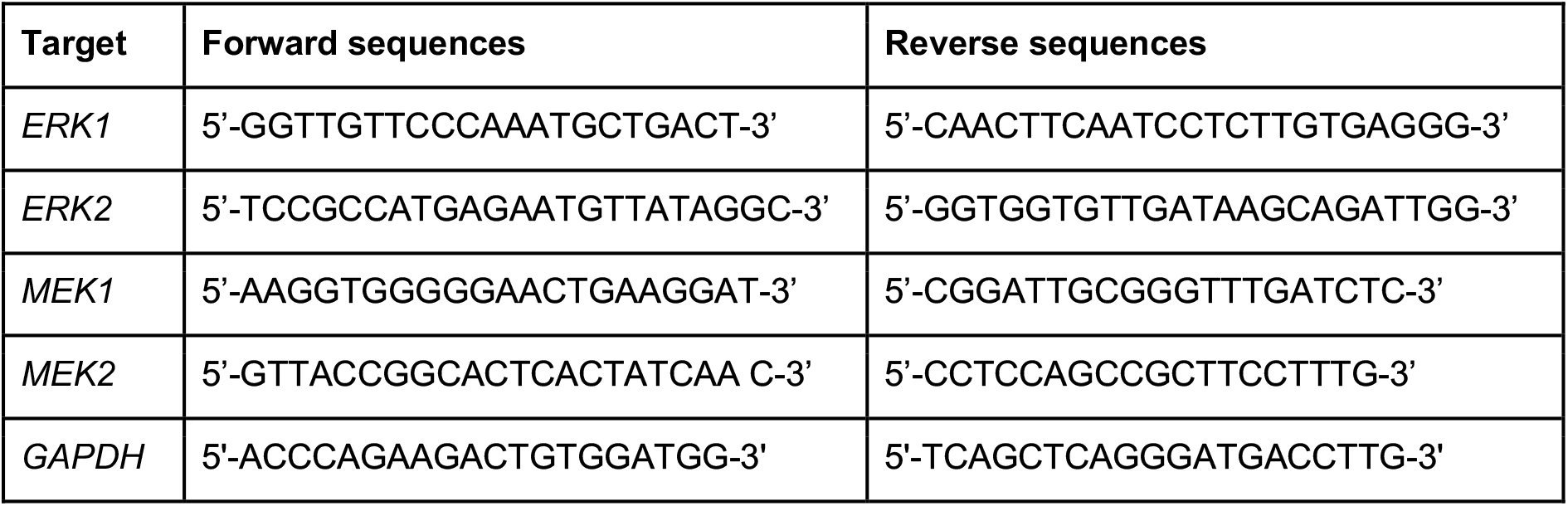

### Immunoblotting

Cells were transfected with 10 nM siPOOL in 6 well plates at 6 × 10^4^ cells/well density and incubated at 37°C with 5% CO_2_ for 72 hours. Cells were lysed in a buffer containing 10 mM Tris HCl, 1 mM EDTA and 1% SDS. Protein concentration was determined with the BCA™ protein assay kit (Thermofisher, 23227). Home cast 10% SDS gels or Novex 4%-20% 10 well Mini Gels (Thermofisher, XP04200) were used for SDS page. Transfer was done using PVDF membranes and a Trans-Blot SD Semi-Dry Electrophoretic Transfer Cell (Bio-Rad). Imaging was done with an Odyssey Fluorescence scanner (Li-COR) (Figure 4C, EV2B). The following primary antibodies were used: anti-total ERK (M7927, Sigma), anti-MEK1 (ab32091, Abcam), anti-MEK2 (ab32517, Abcam), anti-BRAF (sc-5284, Santa Cruz), anti-CRAF (9422S, Cell Signaling Technology), anti-SOS1 (610096, Biosciences), anti-GRB2 (PA5-17692, Invitrogen) and anti-RSK2 (sc-9986, Santa Cruz). Anti-GAPDH (sc-32233, Santa Cruz) or anti-Actin (A2066, Merck) was used as protein of reference. For the secondary antibodies, we used the IRDye 680LT donkey anti-mouse IgG (926-68022, Li-COR), IRDye 800CW goat anti-mouse (926-32210, Li-COR) and IRDye 800CW donkey anti-rabbit (926-32213, Li-COR). Protein quantification was done with the Image Studio™ Lite software.

### Time-series feature extraction

We used custom scripts to extract features of ERK responses to transient optoFGFR input (Figure 2B, EV1A,B), sustained GF input (Figure EV1C,D) and transient optoSOS input (EV4A). The maximum peak (maxPeak) is the absolute value of the highest ERK activity in the trajectory. To estimate the full width at half maximum (FWHM), we first removed the baseline of the trajectories and increased their sampling frequency by a factor 30 with spline interpolation. On the resulting trajectory, we applied a “walk” procedure to quantify the FWHM. In this method, a pointer walks left and right (*i*.*e*. opposite and along the direction of time respectively) from the maximum point of the trajectory. The pointer stops whenever the half maximum value is crossed. Both stops define a left and a right border, the time difference between these 2-border time-points gives the FWHM. To avoid reporting aberrant FWHM values in cases where a peak cannot be clearly defined, we excluded FWHM calculation for trajectories where the fold change between the baseline (mean activity before stimulation) and the maximum value of the trajectory was below a threshold manually defined. ERKpostStim is the absolute value of ERK activity extracted 9 minutes after the last stimulation pulse to evaluate ERK adaptation. Statistical analysis (Figure EV1A,B) was done by comparing all conditions to the 20-minute interval stimulation patterns with a Wilcoxon test using the FDR p-value correction (NS: non-significant, *<0.01, **<0.001, ***<0.0001, ****<0.00001).

To evaluate ERK phenotypes under siRNA perturbations in response to sustained optoFGFR or optoSOS input (Figure 4D, EV3A, 5F, EV4F), we extracted the baseline (average ERK activity on 5 timepoints before stimulation), the maxPeak (maximum ERK activity within a 10-minute time window following the start of the stimulation) and the ERKpostStim (ERK activity at a fixed timepoint post-stimulation (t_optoFGFR_ = 42 min and t_optoSOS_ = 40 min)) from 3 technical replicates. To avoid heterogeneity due to differences in optogenetic expression, we focused our analysis on cells with high optogenetic expression. The obtained baseline, maxPeak and ERKpostStim for each siRNA perturbation was z-scored to the non-targeting siRNA (*CTRL*). Non-significant results were manually set to grey. Statistical analysis was done by comparing each perturbation to the control with a Wilcoxon test using the FDR p-value correction (NS: non-significant, *<0.05, **<0.005, ***<0.0005, ****<0.00005).

For the comparison of both optogenetic systems (Figure 5E, EV4C), ERK baseline was obtained by averaging ERK activity on 5 timepoints before stimulation and ERK maxPeak was extracted within a 10-minute time window following the start of the stimulation. Statistical analysis was done by comparing low and high expressing cells within and across optogenetic systems with a Wilcoxon test using the FDR p-value correction (NS: non-significant, *<0.05, **<0.005, ***<0.0005, ****<0.00005).

To quantify the efficiency of the three MAPK inhibitors on the reduction of ERK amplitudes under sustained high optoFGFR or optoSOS input (Figure 6), extraction of the maxPeak was limited by the fact that several concentrations led to a full suppression of ERK amplitudes. Therefore, we extracted ERK amplitudes at a fixed time point following the start of the stimulation (t_fixed optoFGFR_ = 15 minutes, t_fixed optoSOS_ = 10 minutes). The obtained ERK amplitudes were then plotted for each concentration for a fixed number of cells randomly selected (Figure 6B,F, EV5D). To calculate the EC_50_ of each drug, we normalized the data by setting the mean ERK responses of the non-treated condition to 1 and the mean ERK responses of the maximum concentration to 0. EC_50_ then was calculated by fitting a Hill function to the mean ERK activity of each concentration (Figure 6C,G, EV5E, Appendix Table 5-7). The heterogeneity of ERK amplitude at the fixed time point was evaluated by computing the normalized standard deviation of the extracted ERK activity per condition (Figure 6D,H, EV5F).

### Identification of ERK dynamics phenotypes using CODEX

To investigate ERK dynamics phenotypes to siRNA perturbations, we first trained a convolutional neural network (CNN) to classify input ERK trajectories into any of the siRNA-perturbed conditions (Figure EV3C). For this purpose, we used a CNN architecture composed of 4 1D-convolution layers with 20 kernels of size 5, followed by a convolution layer with 20 kernels of size 3 and one layer of 10 kernels of size 3. The responses are then pooled with global average pooling to generate a vector of 10 features that is passed to a (10,63) fully connected layer for classification. Each convolutional layer is followed by ReLU and batch normalization. The CNN was trained to minimize the cross-entropy loss, with L2 weight penalty of 1e^-3^.

To identify siRNA treatments that induced a distinctive phenotype, we selected the 10 conditions for which the CNN classification precision was the highest on the validation set (Appendix Table S4, “CODEX accuracy”). To these 10 conditions, we also added the negative control (non-targeting siRNA (*CTRL*)). We trained a second CNN, with the same architecture and training parameters, but limited to recognizing the 11 selected treatments to obtain a clear embedding of these hits. With this new model, we extracted the features used for the classification of the trajectories (i.e. the input representation after the last convolution layer) and projected them with tSNE (Python’s *sklearn* implementation, perplexity of 100, learning rate of 600 and 2500 iterations) (Figure EV3D). We selected 10 prototype curves for each treatment by taking the trajectories for which the second CNN’s classification confidence (*i*.*e*. the probability for the actual class of the inputs) were the highest in the validation set (Figure 4E, “CODEX”).

To visualize the ERK dynamics landscape in MCF10A WT cells and in MCF10A cells overexpressing ErbB2, we trained one CNN for each cell line. These CNNs were trained to recognize the drug treatment applied on cells, using single-cell ERK traces as input. The architecture of the CNNs is the same as described previously. The only difference lies in the number of outputs in the final fully connected layer, which were set to the number of drug treatments. Features used for the classification of the trajectories were then projected with tSNE (Figure 7D,H, EV5J,K).

To identify clusters gathering similar ERK dynamics (Figure 7E,F,I,J), we clustered trajectories based on their CNN features using a partition around medoids (PAM). This iterative algorithm is similar to K-means clustering. PAM defines the cluster centers (i.e. the medoids) as the observed data points which minimize the median distances to all other points in its own cluster. This makes PAM more robust to outliers than K-means which uses the average coordinates of a cluster to define its center. Representative trajectories were obtained by taking the medoids of each cluster and their four closest neighbors (Figure 7G,K). Distances between points were defined with the Manhattan distance between the scaled CNN features (zero mean and unit variance). We manually verified that these clusters captured an actual trend by visualizing trajectories in each cluster with the interactive CODEX application.

### Peak detection and classification of oscillatory trajectories

The number of ERK activity peaks was calculated with a custom algorithm that detects local maxima in time series. First, we applied a short median filter to smoothen the data with a window width of 3 time points. Then, we ran a long median filter to estimate the long-term bias with a window width of 15 time points. This bias was then subtracted from the smoothed time series and we only kept the positive values. If no point in this processed trajectory was exceeding a manual threshold of 0.075, all variations were considered as noise and no peak was extracted from the trajectory. The remaining trajectories were then rescaled to [0,1]. Finally, peaks were detected as points that exceeded a threshold which was manually set to 0.1. Peaks that were found before the first stimulation or after the last stimulation were filtered out.

The classification of trajectories into oscillatory and non-oscillatory behaviors was performed after the peak detection step. Cells were called oscillatory if at least 3 peaks were detected with the peak detection procedure (Figure 4F). Statistical analysis was done using a pairwise t-test comparing each perturbation to the control for high and low levels of optoFGFR independently, with FDR p-value correction (*<0.05, **<0.005, ***<0.0005, ****<0.00005).

### Data availability

The datasets used in this study as well as all R codes used for further analysis are available at https://data.mendeley.com/datasets/st36dd7k23/1. Source code for the inference algorithm, model files and results are available at https://github.com/Mijan/LFNS_optoFGFR.

## Acknowledgements

This work was supported by SystemsX.ch, Swiss Cancer League and Swiss National Science Foundation grants to Olivier Pertz, by the H2020-MSCA-IF, project No. 89631 - NOSCAR to Agne Frismantiene and by the European Union’s Horizon 2020 and innovation program under grant agreement No. 730964 (TRANSVAC project) to Mustafa Khammash. We thank Won Do Heo for sharing the optoFGFR plasmid, Kazuhiro Aoki for sharing the pPBbSr2-MCS plasmid, and David Hacker for sharing the pSB-HPB plasmid. We thank the Microscopy Imaging Center of the University of Bern for its support.

## Authors contribution

O.P. and C.D. designed the study. C.D. developed the optogenetic systems and imaging pipelines. CD performed the experiment and image analysis on NIH3T3. A.F and P.A.G. performed the experiments and image analysis on MCF10A. M.D. developed the processing pipelines. C.D processed the data. C.D., M.-A.J., A.F and P.A.G. analyzed the data. M.-A.J. conducted the CNN analysis. J.M. performed mathematical modeling. O.P and M.K. supervised the work. O.P., C.D. and J.M. wrote the paper.

