## Supplementary for "Optogenetic actuator/ERK biosensor circuits identify MAPK network nodes that shape ERK dynamics"

### Expanded View Figures

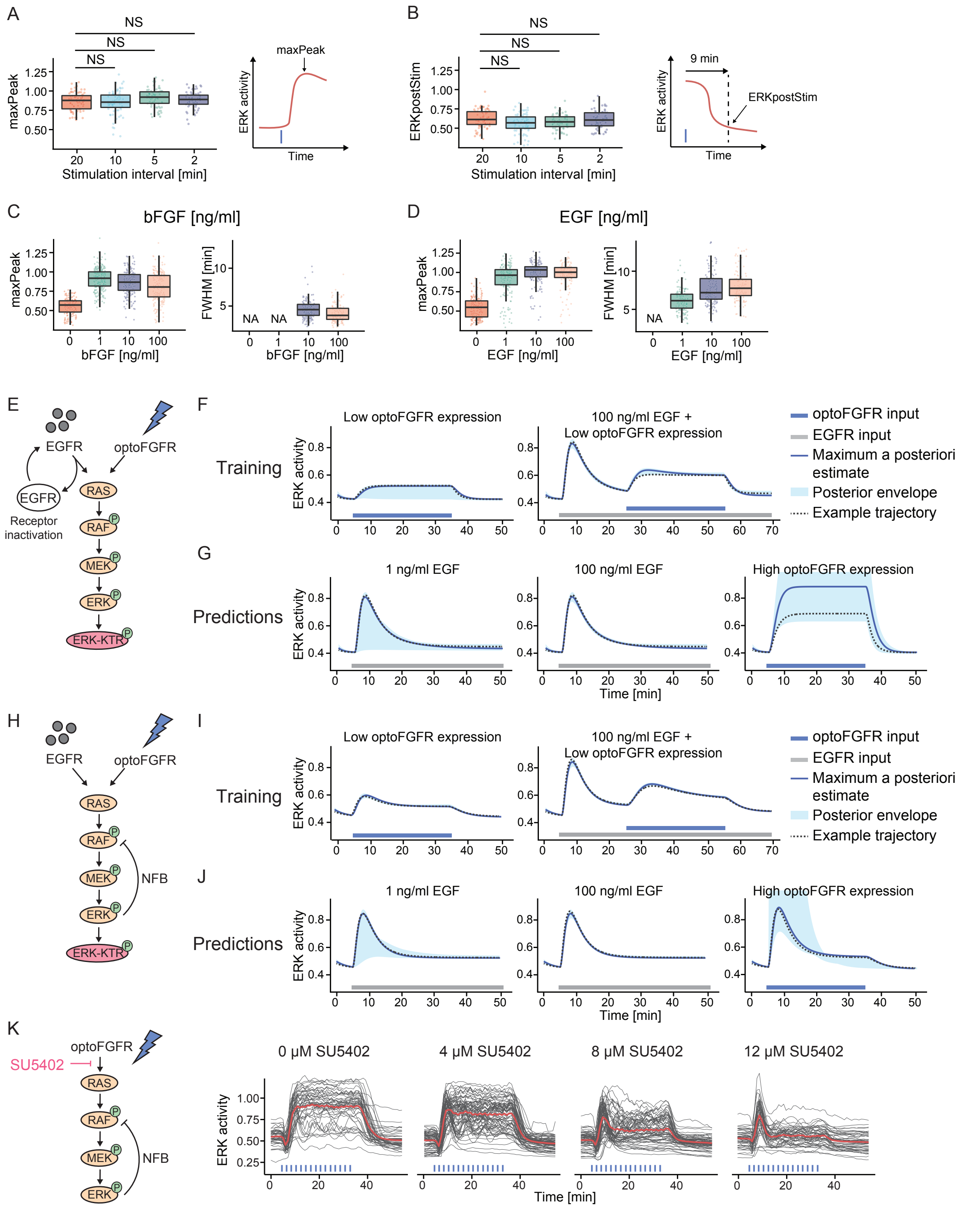

**Figure EV1: ERK dynamics evoked by optoFGFR versus endogenous RTKs highlight different MAPK regulatory mechanisms. (A-B)** MaxPeak and ERKpostStim quantifications of ERK responses for each stimulation pattern shown in Figure 2C ($N_{min}$ = 63 cells per condition). MaxPeak was quantified within a 10 min time window following the first stimulation pulse. ERKpostStim was extracted 9 minutes after the last pulse. Statistical analysis was done using a Wilcoxon test comparing each condition to the 20-minute interval pattern (NS: non-significant, *<0.01, **<0.001, ***<0.0001, ****<0.00001, FDR p-value correction method). **(C-D)** MaxPeak and FWHM quantification of ERK responses shown in Figure 3A ($N_{min}$ = 150 cells per condition) **(C)** and in Figure 3B ($N_{min}$ = 130 cells per condition) **(D)**. **(E-J)** Alternative mathematical models without the presence of a NFB **(E-G)** or without EGFR regulation **(H-J)** were trained on the average ERK responses shown in Figure 3E and used to predict ERK responses to the different light and EGF input. The maximum a posteriori (MAP) estimate, the posterior envelope indicating the predictive density of the estimation as well as one example trajectory are shown. **(K)** ERK responses to sustained optoFGFR input (D = 18 mJ/cm$^2$) under dose response inhibition with the FGFR inhibitor (SU5402).

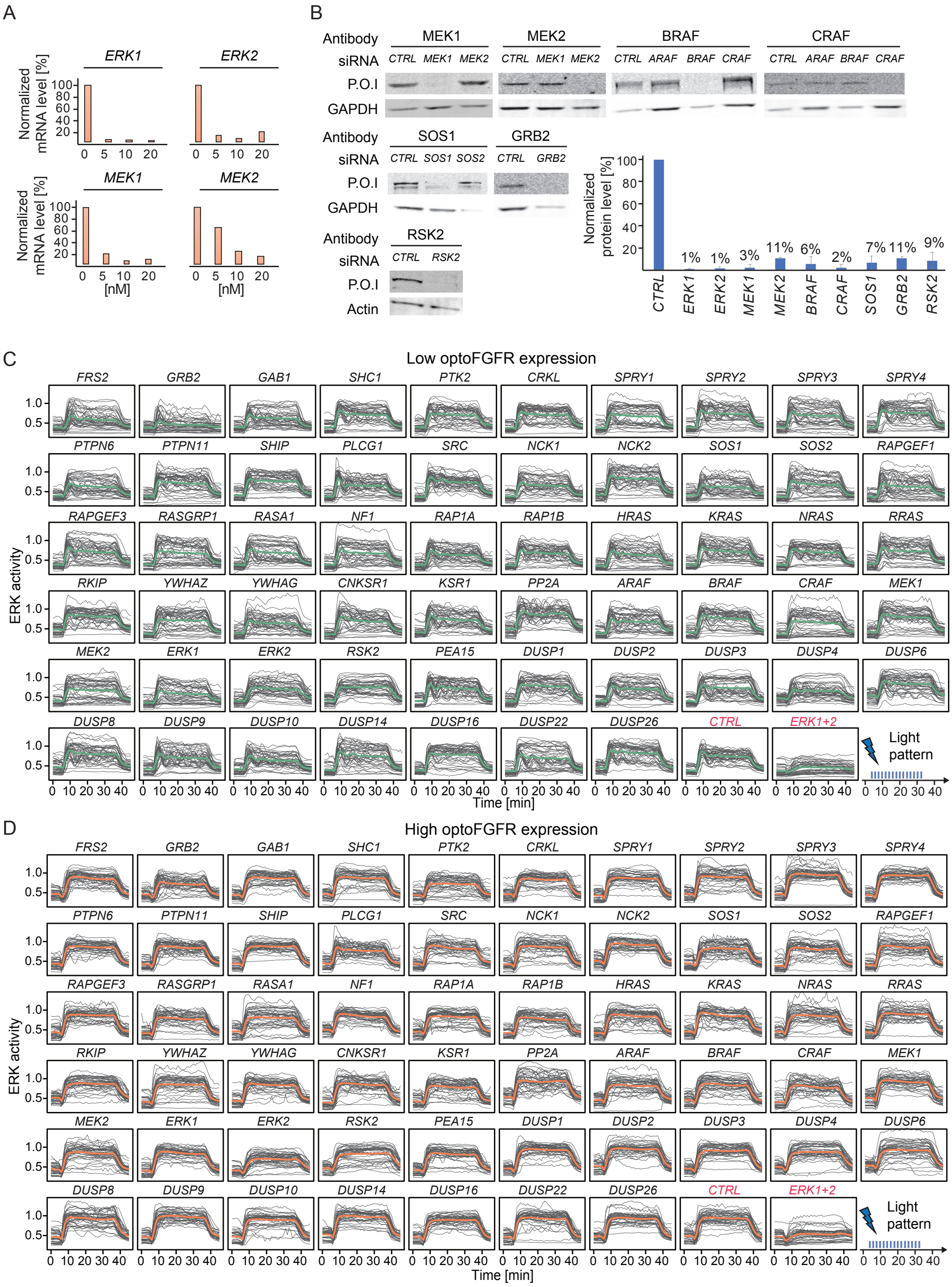

**Figure EV2: RNA interference screen reveals that ERK dynamics remain unaffected in response to perturbation of most MAPK signaling nodes. (A)** mRNA level quantification (RT-qPCR) of cells transfected with different concentrations of siRNA targeting *ERK1, ERK2, MEK1* and *MEK2*. Data were normalized to the amount of *GAPDH* mRNA and relatively expressed to the corresponding mRNA in non-transfected cells. **(B)** Western blot analysis of cells transfected with 10 nM of siRNA against the different *MEKs* and *RAFs* isoforms, *SOS1*, *GRB2 and RSK2*. Western blot analysis of cells transfected with the *ERKs* isoforms is shown in Figure 4C. Remaining protein levels were quantified by normalizing the amount of protein of interest (P.O.I) to the amount of GAPDH or actin protein and shown relatively to the corresponding protein level in the CTRL (N = 2 replicates, except for *SOS1*). **(C-D)** Single-cell ERK responses under the different RNAi perturbations (sustained optoFGFR input, D = 18 mJ/cm$^2$, N

= 40 cells randomly selected from low **(C)** and high **(D)** optoFGFR expression, out of at least 212 cells per perturbation from 3 technical replicates).

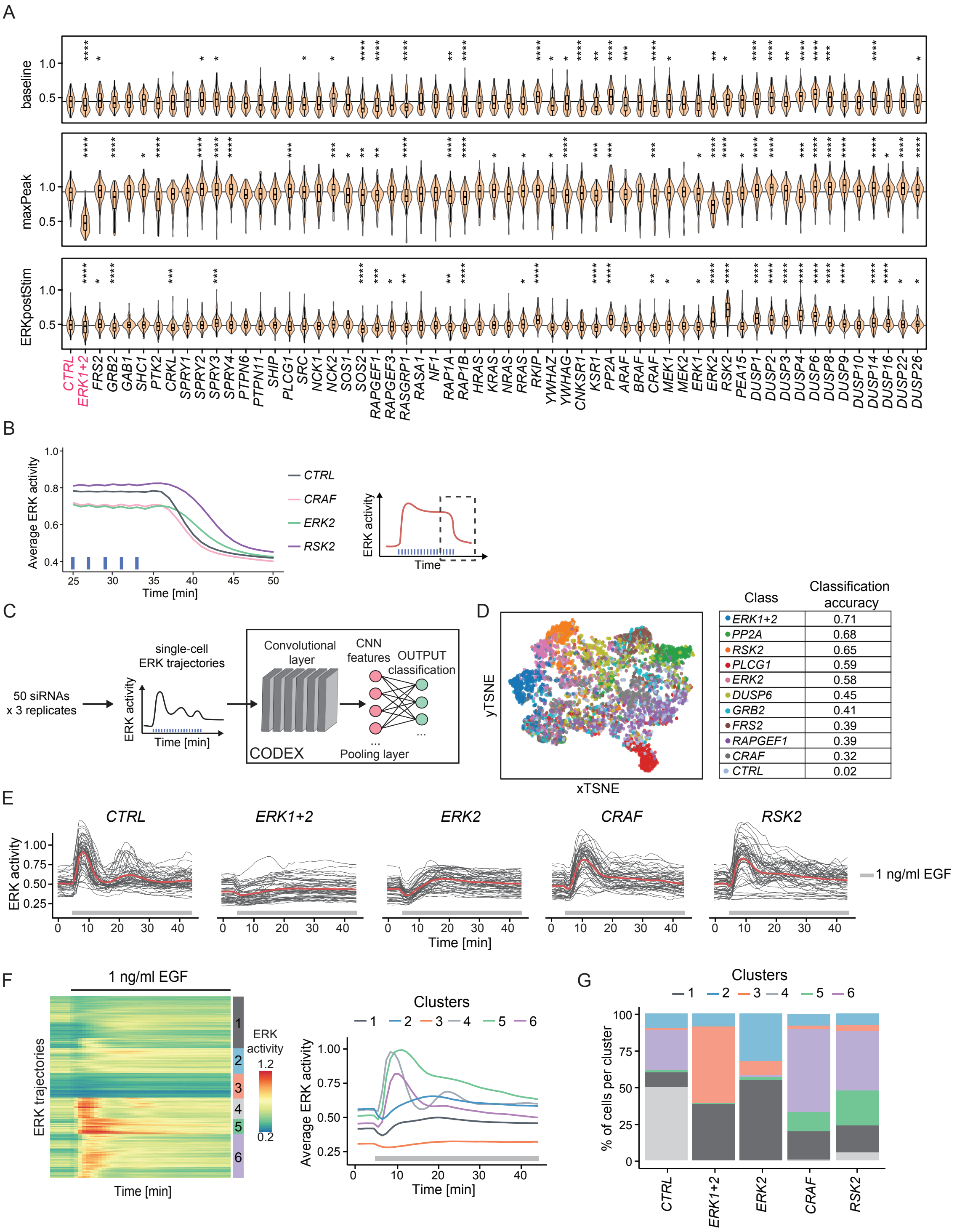

**Figure EV3: Investigation of penetrant siRNA ERK dynamic phenotypes. (A)** Violin plot distributions of the baseline, maxPeak and ERKpostStim of single-cell ERK responses under sustained high optoFGFR input (D = 18 mJ/cm$^2$, $N_{min}$ = 126 cells with high optoFGFR expression per treatment, from 3 technical replicates). Statistical analysis was done using a Wilcoxon test comparing each treatment to the control (*<0.05, **<0.005, ***<0.0005, ****<0.00005, FDR p-value correction method). **(B)** Average ERK response for selected siPOOLs affecting ERK adaptation (ERKpostStim in Figure 4D) (sustained optoFGFR input, D = 18 mJ/cm$^2$, $N_{min}$ = 400 cells per condition from 3 technical replicates). **(C)** CODEX was trained to recognize single-cell ERK responses evoked by sustained optoFGFR input (D = 18 mJ/cm$^2$) under the different RNAi perturbations ($N_{min}$ = 212 cells per perturbation from 3 technical replicates). **(D)** tSNE projection of the CNN features from CODEX trained on the 10 perturbations for which the classification accuracy on the validation set was the highest when trained on all perturbations (Appendix Table S4, see Material and Methods), together with the non-targeting siRNA (*CTRL*). **(E)** Single-cell ERK trajectories under 1 ng/ml sustained EGF input (added at t = 5 minutes) for selected RNAi perturbations (N = 50 cells per condition). **(F)** Hierarchical clustering (Euclidean distance and Ward linkage) of single-cell ERK trajectories shown in (E) ($N_{CTRL}$ = 300 cells, $N_{ERK1+2}$ = 270 cells, $N_{ERK2}$ = 320 cells, $N_{CRAF}$ = 240 cells, $N_{RSK2}$ = 340 cells). The number of clusters was empirically defined to resolve the different ERK dynamics. Average ERK responses per cluster are displayed on the right. **(G)** Proportion of single-cell ERK trajectories per cluster shown in (F).

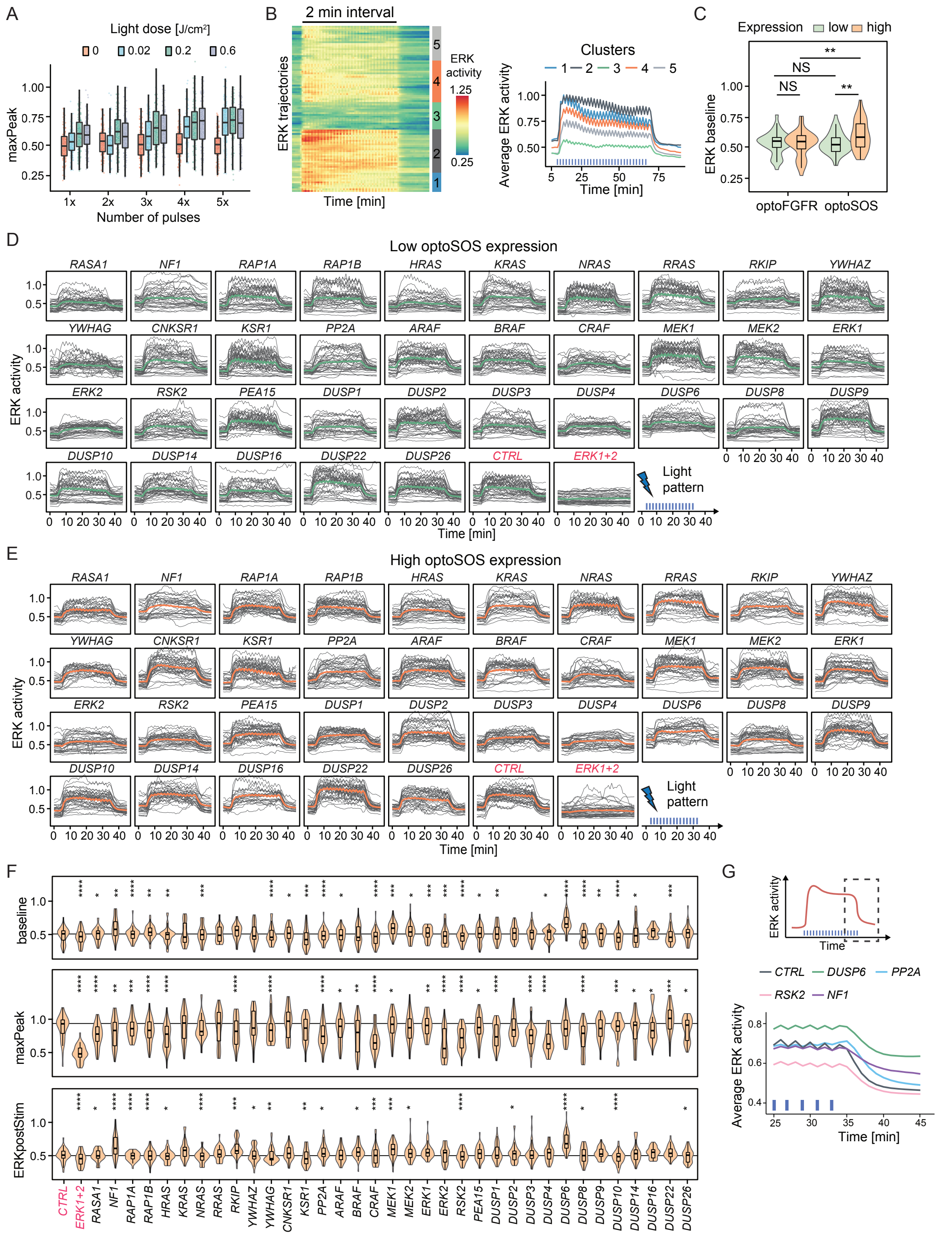

**Figure EV4: Direct optogenetic activation of RAS highlights different ERK dynamics phenotypes than optoFGFR input. (A)** Quantification of the maxPeak of ERK dynamics evoked by transient optoSOS input using different light doses (color code) and different numbers of 100 ms pulses (x-axis) repeated every 20 seconds. **(B)** Hierarchical clustering (Maximum distance and Ward D2 linkage) of ERK responses under 2-minute interval optoSOS input shown in Figure 5C (D = 0.6 J/cm$^2$, N = 90 cells). The number of clusters was empirically defined to resolve the different ERK amplitudes. Average ERK responses per cluster are displayed on the right. **(C)** Quantification of the baseline of single-cell

ERK responses under sustained optoFGFR (Figure 2F, D = 18 $mJ/cm^2$) and optoSOS (Figure 5D, D = 0.6 $J/cm^2$) input for low or high expression of each optogenetic system (N = 40 cells per condition). Statistical analysis was done using a Wilcoxon test, comparing each condition to each other ($N_{min}$ = 48 cells per condition, NS: non-significant, *<0.05, **<0.005, ***<0.0005, ****<0.00005, FDR p-value correction method). **(D-E)** ERK responses under RNAi perturbations targeting MAPK signaling nodes active below RAS (sustained optoSOS input, D = 0.6 $J/cm^2$, N = 40 cells from low **(D)** and high **(E)** optoSOS expressing cells for each perturbation, randomly selected out of at least 193 trajectories from 3 technical replicates). **(F)** Violin plot distributions of the baseline, maxPeak and ERKpostStim of single-cell ERK responses under sustained high optoSOS input (D = 0.6 $J/cm^2$, $N_{min}$ = 33 cells with high optoSOS expression per treatment, from 3 technical replicates). Statistical analysis was done using a Wilcoxon test comparing each treatment to the control (*<0.05, **<0.005, ***<0.0005, ****<0.00005, FDR p-value correction method). **(G)** Average ERK response for selected siPOOLs affecting ERK adaptation (ERKpostStim in Figure 5F) (sustained optoSOS input, D = 0.6 $J/cm^2$, $N_{min}$ = 270 cells per condition from 3 technical replicates).

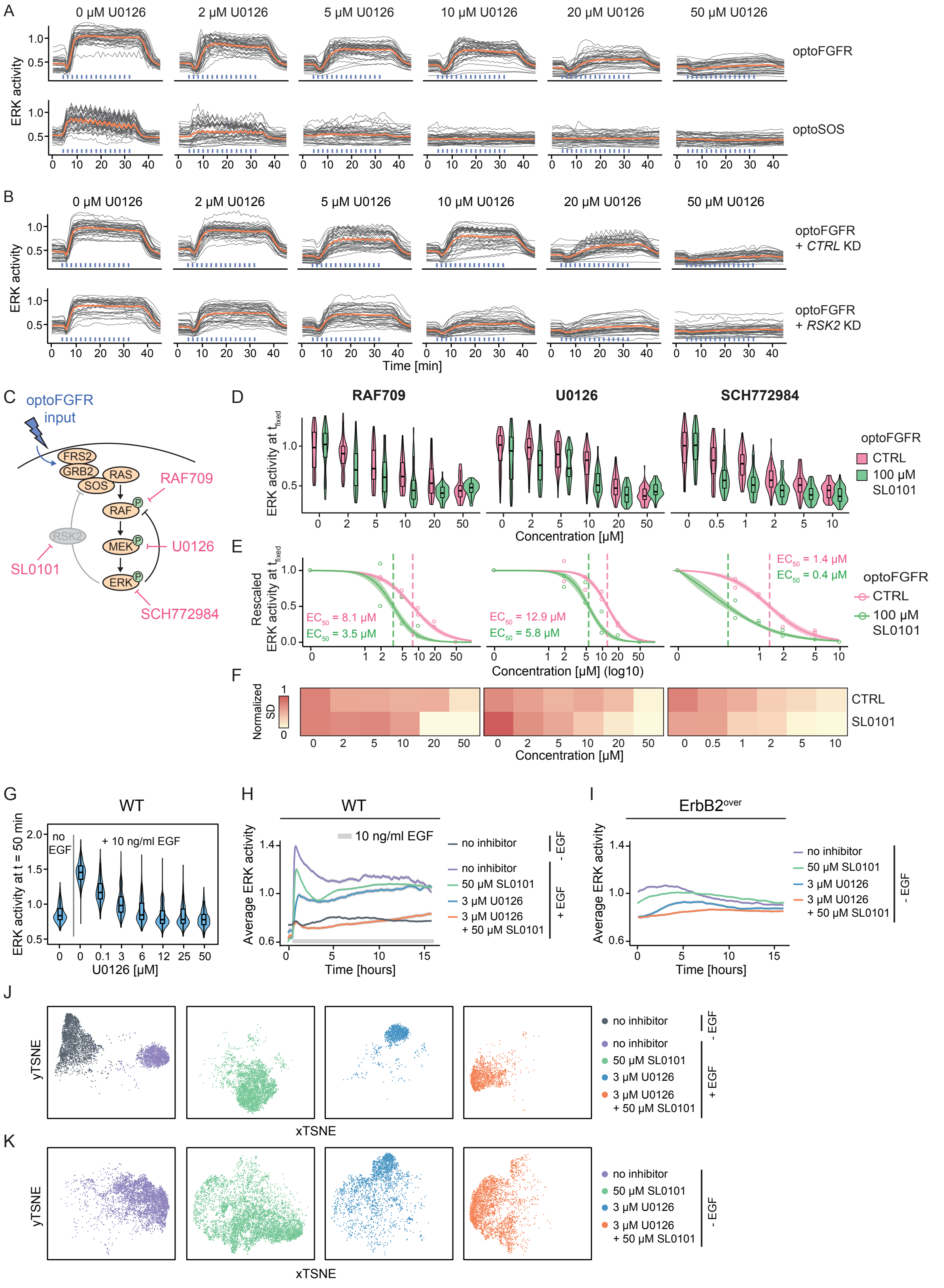

**Figure EV5: Perturbation of the RSK2-mediated NFB increases the efficiency of RAS, MEK and ERK targeting drugs. (A)** Single-cell ERK trajectories from high optoFGFR or high optoSOS expressing cells treated with U0126 dose response (N = 40 cells per condition, randomly selected out of at least 200 cells from 3 technical replicates, sustained optoFGFR (D = 18 mJ/cm$^2$) or optoSOS (D =

0.6 J/cm$^2$) input). **(B)** Single-cell ERK trajectories from high optoFGFR expressing cells treated with U0126 dose response for *CTRL* or *RSK2* KD cells (N = 40 cells per condition (apart from *RSK2* KD + 0 µM U0126 (32 cells), from 2 technical replicates for *RSK2* KD and 1 replicate for *CTRL* KD, sustained optoFGFR input (D = 18 mJ/cm$^2$)). For (A) and (B), data for RAF709 and SCH772984 are available in supplementary material. **(C)** Schematic representation of the optoFGFR system untreated (RSK2-mediated feedback dependent) or treated with an RSK2 inhibitor SL0101 (RSK2-mediated feedback independent) targeted with the B/CRAF (RAF709), the MEK (U0126) or the ERK (SCH772984) inhibitor. **(D)** Single-cell ERK amplitudes from sustained high optoFGFR input (D = 18 mJ/cm$^2$) under different concentrations of the MAPK inhibitors, extracted at a fixed time point ($t_{fixed\ optoFGFR}$ = 15 minutes, N = 70 cells with high optoFGFR expression per condition (except for 100 µM SL0101 + 0 µM U0126 (36 cells) randomly selected from 2 technical replicates). **(E)** A Hill function was fit to the normalized mean ERK activity as shown in (D) ($N_{min}$ = 36 cells per condition). Shaded area indicates the 95% CI and dashed lines the $EC_{50}$. **(F)** Normalized standard deviation of ERK amplitudes shown in (D) ($N_{min}$ = 36 cells per condition). **(G)** U0126 dose response analysis in MCF10A WT cells stimulated with 10 ng/ml EGF at t = 30 minutes. ERK responses were extracted at a fixed time point following the steep rising phase ($t_{fixed}$ = 50 minutes, N = 140 cells per condition). **(H-I)** Average ERK responses of MCF10A WT cells with EGF stimulation (10 ng/ml EGF added at t = 30 minutes) **(H)** or of ErbB2 overexpressing ($ErbB2^{over}$) MCF10A cells without EGF stimulation **(I)** under no inhibitor, RSK (SL0101) inhibitor, MEK (U0126) inhibitor or a combination of both. Shaded areas indicate the 95% CI. **(J-K)** tSNE projection of CODEX's CNN features per treatment from ERK activity of MCF10A WT cells **(J)** or MCF10A $ErbB2^{over}$ cells **(K)** treated with 50 µM SL0101, 3 µM U0126 or a combination of both.

### Appendix Figures

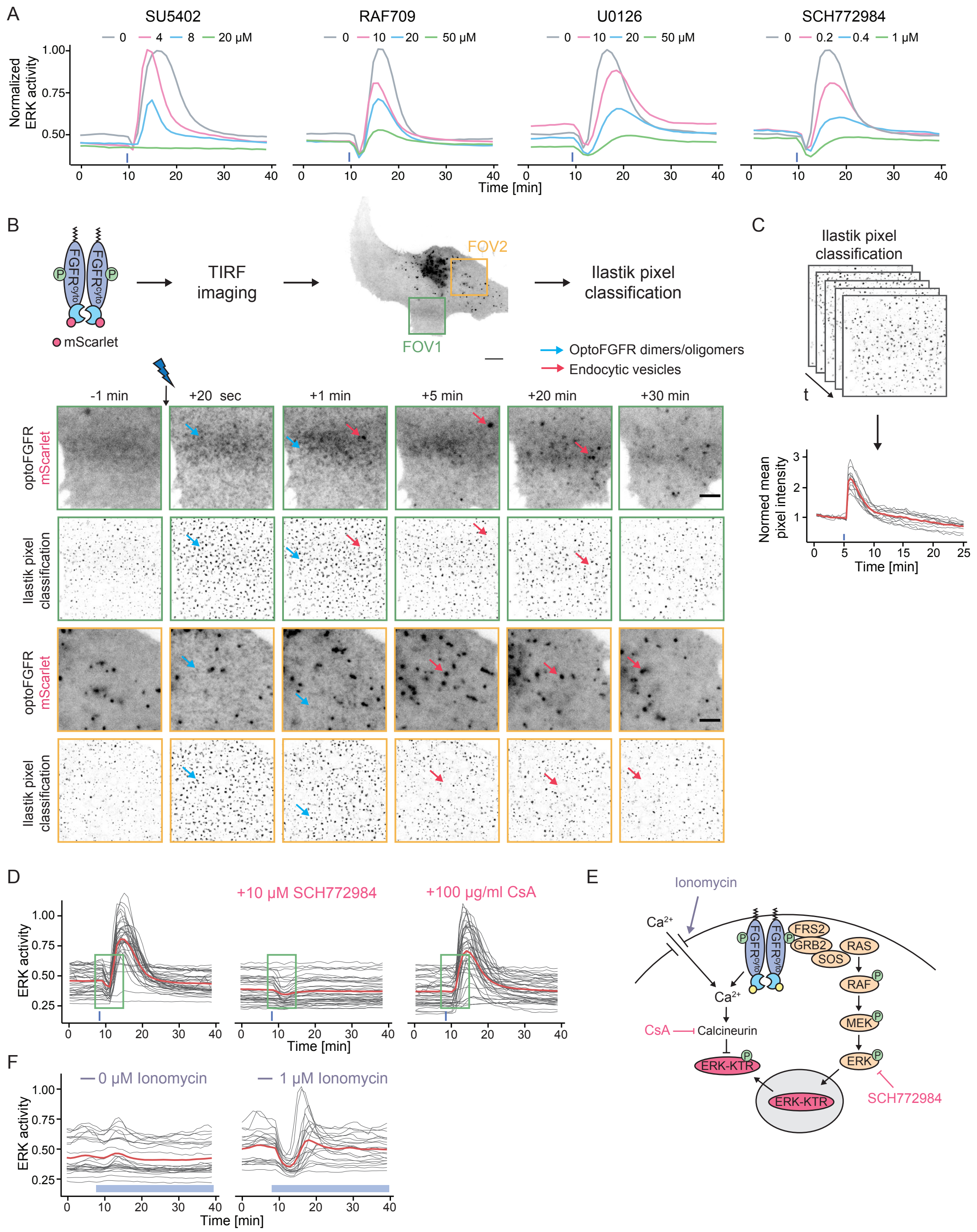

**Appendix Figure S1: An optogenetic actuator-biosensor genetic circuit to study input-dependent ERK dynamics. (A)** Average ERK responses of cells stimulated with a transient optoFGFR input (D = 18 mJ/cm$^2$) under increasing concentrations of FGFR (SU5402), B/CRAF (RAF709), MEK (U0126) and ERK (SCH772984) inhibitors. ERK responses were normalized to the 0 µM condition for each drug ($N_{min}$ = 40 cells per condition). **(B-C)** Quantification of optoFGFR dimerization in response to a transient 470 nm light input using a mScarlet tagged version of optoFGFR. **(B)** Cells expressing optoFGFR-mScarlet were imaged every 20 seconds with a 100x TIRF objective. Segmentation of optoFGFR dimers/oligomers was performed with the pixel classification module of Ilastik to specifically segment optoFGFR dimer-/oligomerization events (blue arrows) and exclude the endocytic vesicles (red arrows). Scale bar: 5 µm (cell) and 2 µm (FOVs). **(C)** OptoFGFR dimer-/oligomerization dynamics was quantified by computing the mean of pixel intensities from the binarized mask obtained with Ilastik. Single-cell trajectories were normalized to the baseline before stimulation (N = 12). **(D)** Investigation of the "dip" (green rectangle) observed in ERK trajectories in response to a transient optoFGFR input. The dip was not suppressed by ERK inhibition (10 µM SCH772984) but was abolished under Calcineurin inhibition (100 µg/ml Cyclosporin A (CsA)). **(E)** Schematized representation of MAPK and $Ca^{2+}$ signaling effect on the ERK-KTR reporter dynamics. **(F)** Validation of $Ca^{2+}$ signaling effect on ERK-KTR by chemically increasing the intracellular $Ca^{2+}$ concentration with 1 µM Ionomycin.

### Appendix Tables

| Species | Notation | Initial Value |
|---|---|---|
| RAS | $RAS$ | 1 |
| Phosphorylated RAS | $RAS^*$ | 0 |
| RAF | $RAF$ | 1 |
| Phosphorylated RAF | $RAF^*$ | 0 |
| MEK | $MEK$ | 1 |
| Phosphorylated MEK | $MEK^*$ | 0 |
| ERK | $ERK$ | 1 |
| Phosphorylated ERK | $ERK^*$ | 0 |
| EGF receptor | $EGFR$ | 1 |
| Active EGF receptor | $EGFR^*$ | 0 |
| Endocytosed EGF receptor | $EGFR_{endo}$ | 0 |
| Negative feedback species | $NFB$ | 1 |
| Active negative feedback species | $NFB^*$ | 0 |
| nuclear KTR | $KTR$ | $1 - ktr_{init}$ |
| cytosolic KTR | $KTR^*$ | $ktr_{init}$ |
| EGF input | $EGF$ | Input variable |
| Light input | $light$ | Input variable |

**Appendix Table S1: Model species, notations, and initial values**

| Nbr | Model Equations |
|---|---|
| 1 | $\dot{EGFR} = -r_{1,2}EGFEGFR + r_{2,1}EGFR^{*} + r_{3,1}EGFR_{endo}$ |
| 2 | $\dot{EGFR}^{*} = r_{1,2}EGFEGFR - (r_{2,1} + r_{2,3})EGFR^{*}$ |
| 3 | $\dot{EGFR}_{endo} = r_{2,3}EGFR^{*} - r_{3,1}EGFR_{endo}$ |
| 4 | $\dot{RAS} = -(k_{1,2}EGFR^{*} + light)\frac{RAS}{K_{1,2} + RAS} + k_{2,1}\frac{RAS^{*}}{K_{2,1} + RAS^{*}}$ |
| 5 | $\dot{RAS}^{*} = (k_{1,2}EGFR^{*} + light)\frac{RAS}{K_{1,2} + RAS} - k_{2,1}\frac{RAS^{*}}{K_{2,1} + RAS^{*}}$ |
| 6 | $\dot{RAF} = -k_{3,4}Ras^{*}\frac{Raf}{K_{3,4} + Raf} + (k_{nfb}NFB^{*} + k_{4,3})\frac{Raf^{*}}{K_{4,3} + Raf^{*}}$ |
| 7 | $\dot{RAF}^{*} = k_{3,4}Ras^{*}\frac{Raf}{K_{3,4} + Raf} - (k_{nfb}NFB^{*} + k_{4,3})\frac{Raf^{*}}{K_{4,3} + Raf^{*}}$ |
| 8 | $\dot{MEK} = -k_{5,6} \cdot RAF^{*}\frac{MEK}{K_{5,6} + MEK} + k_{6,5}\frac{MEK^{*}}{K_{6,5} + MEK^{*}}$ |
| 9 | $\dot{MEK}^{*} = k_{5,6} \cdot RAF^{*}\frac{MEK}{K_{5,6} + MEK} - k_{6,5}\frac{MEK^{*}}{K_{6,5} + MEK^{*}}$ |
| 10 | $\dot{ERK} = -k_{7,8} \cdot MEK^{*}\frac{ERK}{K_{7,8} + ERK} + k_{8,7}\frac{ERK^{*}}{K_{8,7} + ERK^{*}}$ |
| 11 | $\dot{ERK}^{*} = k_{7,8} \cdot MEK^{*}\frac{ERK}{K_{7,8} + ERK} - k_{8,7}\frac{ERK^{*}}{K_{8,7} + ERK^{*}}$ |
| 12 | $\dot{NFB} = -f_{1,2} \cdot ERK^{*}\frac{NFB}{F_{1,2} + NFB} + f_{2,1}\frac{NFB^{*}}{F_{2,1} + NFB^{*}}$ |
| 13 | $\dot{NFB}^{*} = f_{1,2} \cdot ERK^{*}\frac{NFB}{F_{1,2} + NFB} - f_{2,1}\frac{NFB^{*}}{F_{2,1} + NFB^{*}}$ |
| 14 | $\dot{KTR} = -\left(k_{9,10} \cdot ERK^{*}\frac{KTR}{K_{9,10} + KTR} + s_{1,2}KTR\right) + s_{2,1}KTR^{*}$ |
| 15 | $\dot{KTR}^{*} = \left(k_{9,10} \cdot ERK^{*}\frac{KTR}{K_{9,10} + KTR} + s_{1,2}KTR\right) - s_{2,1}KTR^{*}$ |

**Appendix Table S2: Model equations**

| Nbr | Parameter Name | Description |
|---|---|---|
| 1 | $r_{1,2}$ | EGF dependent EGFR activation rate |
| 2 | $r_{2,3}$ | EGFR endocytosis rate |
| 3 | $r_{3,1}$ | EGFR recycling rate |
| 4 | $r_{2,1}$ | EGFR deactivation rate |
| 5 | $k_{1,2}$ | EGFR dependent RAS phosphorylation rate |
| 6 | $K_{1,2}$ | Michaelis constant RAS phosphorylation |
| 7 | $k_{2,1}$ | RAS dephosphorylation rate |
| 8 | $K_{2,1}$ | Michaelis constant RAS dephosphorylation |
| 9 | $k_{3,4}$ | RAF phosphorylation rate |
| 10 | $K_{3,4}$ | Michaelis constant RAF phosphorylation |
| 11 | $k_{nfb}$ | NFB effect on RAF dephosphorylation |
| 12 | $k_{4,3}$ | RAF dephosphorylation rate |
| 13 | $K_{4,3}$ | Michaelis constant RAF dephosphorylation |
| 14 | $k_{nfb}$ | NFB effect on RAF dephosphorylation |
| 15 | $k_{5,6}$ | RAF dependent MEK phosphorylation rate |
| 16 | $K_{5,6}$ | Michaelis constant MEK phosphorylation |
| 17 | $k_{6,5}$ | MEK dephosphorylation rate |
| 18 | $K_{6,5}$ | Michaelis constant MEK dephosphorylation |
| 19 | $k_{7,8}$ | MEK dependent ERK phosphorylation rate |
| 20 | $K_{7,8}$ | Michaelis constant ERK phosphorylation |
| 21 | $k_{8,7}$ | ERK dephosphorylation rate |
| 22 | $K_{8,7}$ | Michaelis constant ERK dephosphorylation |
| 23 | $k_{9,10}$ | ERK dependent KTR translocation rate |
| 24 | $K_{9,10}$ | Michaelis constant KTR translocation |
| 25 | $s_{1,2}$ | ERK independent KTR translocation rate |
| 26 | $s_{2,1}$ | nuclear KTR translocation rate |
| 27 | $f_{1,2}$ | ERK dependent NFB activation rate |
| 24 | $F_{1,2}$ | Michaelis constant NFB activation |
| 25 | $f_{2,1}$ | NFB deactivation rate |
| 26 | $F_{2,1}$ | Michaelis constant NFB deactivation |
| 27 | $ktr_{init}$ | Fraction of initial cytosolic KTR |
| 27 | $\sigma$ | Standard deviation of the measurement noise |

**Appendix Table S3: Model parameters**

| siRNA name | Gene name | Mouse gene ID | CODEX accuracy |
|---|---|---|---|
| ***FRS2*** | Frs2 | 327826 | 0.385 |
| ***GRB2*** | Grb2 | 14784 | 0.414 |
| ***GAB1*** | Gab1 | 14388 | 0.082 |
| ***SHC1*** | Shc1 | 20416 | 0.217 |
| ***PTK2*** | Ptk2 | 14083 | 0 |
| ***CRKL*** | Crkl | 12929 | 0.015 |
| ***SPRY1*** | Spry1 | 24063 | 0.113 |
| ***SPRY2*** | Spry2 | 24064 | 0.058 |
| ***SPRY3*** | Spry3 | 236576 | 0.174 |
| ***SPRY4*** | Spry4 | 24066 | 0.083 |
| ***PTPN6*** | Ptpn6 | 15170 | 0.11 |
| ***PTPN11*** | Ptpn11 | 19247 | 0.128 |
| ***SHIP*** | Inpp5d | 16331 | 0.125 |
| ***PLCG1*** | Plcg1 | 18803 | 0.589 |
| ***SRC*** | Src | 20779 | 0.136 |
| ***NCK1*** | Nck1 | 17973 | 0.002 |
| ***NCK2*** | Nck2 | 17974 | 0.069 |
| ***SOS1*** | Sos1 | 20662 | 0.319 |
| ***SOS2*** | Sos2 | 20663 | 0.041 |
| ***RAPGEF1*** | Rapgef1 | 107746 | 0.385 |
| ***RAPGEF3*** | Rapgef3 | 223864 | 0.052 |
| ***RASGRP1*** | Rasgrp1 | 19419 | 0.128 |
| ***RASA1*** | Rasa1 | 218397 | 0 |
| ***NF1*** | Nf1 | 18015 | 0.112 |
| ***RAP1A*** | Rap1a | 109905 | 0.024 |
| ***RAP1B*** | Rap1b | 215449 | 0.099 |
| ***HRAS*** | Hras | 15461 | 0.004 |
| ***KRAS*** | Kras | 16653 | 0.126 |
| ***NRAS*** | Nras | 18176 | 0.002 |
| ***RRAS*** | Rras | 20130 | 0.145 |
| ***RKIP*** | Pebp1 | 23980 | 0.045 |
| ***YWHAZ*** | Ywhaz | 22631 | 0.044 |
| ***YWHAG*** | Ywhag | 22628 | 0.043 |
| ***CNKSR1*** | Cnksr1 | 194231 | 0.115 |
| ***KSR1*** | Ksr1 | 16706 | 0.085 |
| ***PP2A*** | Ppp2ca | 19052 | 0.682 |
| ***ARAF*** | Araf | 11836 | 0.021 |
| ***BRAF*** | Braf | 109880 | 0.031 |
| ***CRAF*** | Raf1 | 110157 | 0.323 |
| ***MEK1*** | Map2k1 | 26395 | 0.056 |
| ***MEK2*** | Map2k2 | 26396 | 0.054 |
| ***ERK1*** | Mapk3 | 26417 | 0.032 |
| ***ERK2*** | Mapk1 | 26413 | 0.578 |
| ***RSK2*** | Rps6ka3 | 110651 | 0.645 |
| ***PEA15*** | Pea15a | 18611 | 0.11 |
| ***DUSP1*** | Dusp1 | 19252 | 0.188 |
| ***DUSP2*** | Dusp2 | 13537 | 0.031 |
| ***DUSP3*** | Dusp3 | 72349 | 0.285 |
| ***DUSP4*** | Dusp4 | 319520 | 0.113 |
| ***DUSP6*** | Dusp6 | 67603 | 0.453 |
| ***DUSP8*** | Dusp8 | 18218 | 0.068 |
| ***DUSP9*** | Dusp9 | 75590 | 0.164 |
| ***DUSP10*** | Dusp10 | 63953 | 0.266 |
| ***DUSP14*** | Dusp14 | 56405 | 0.066 |
| ***DUSP16*** | Dusp16 | 70686 | 0.035 |
| ***DUSP22*** | Dusp22 | 105352 | 0.012 |
| ***DUSP26*** | Dusp26 | 66959 | 0.155 |
| ***CTRL*** | - | - | 0.021 |
| ***ERK1+2*** | - | - | 0.711 |

**Appendix Table S4: Names of signaling nodes targeted with RNA interference and CODEX classification accuracy**

| System | Inhibitor | $EC_{50}$ | Lower | Upper |
|---|---|---|---|---|
| optoFGFR | RAF709 | 7.9316 | 7.5452 | 8.3180 |
| | U0126 | 10.3015 | 9.8784 | 10.7245 |
| | SCH772984 | 1.1024 | 1.0456 | 1.1592 |
| optoSOS | RAF709 | 4.0766 | 3.7665 | 4.3866 |
| | U0126 | 1.6256 | 1.4082 | 1.8430 |
| | SCH772984 | 0.2163 | 0.1610 | 0.2717 |

**Appendix Table S5: $EC_{50}$ with upper and lower values for the fit shown in Figure 6C**

| System | Inhibitor | $EC_{50}$ | Lower | Upper |
|---|---|---|---|---|
| optoFGFR + *CTRL* KD | RAF709 | 8.2045 | 7.5718 | 8.8373 |
| | U0126 | 12.7216 | 11.9047 | 13.5385 |
| | SCH772984 | 1.4052 | 1.3096 | 1.5008 |
| optoFGFR + *RSK2* KD | RAF709 | 4.0463 | 3.6547 | 4.4380 |
| | U0126 | 5.2965 | 4.7404 | 5.8526 |
| | SCH772984 | 0.6369 | 0.5738 | 0.6999 |

**Appendix Table S6: $EC_{50}$ with upper and lower values for the fit shown in Figure 6G**

| System | Inhibitor | $EC_{50}$ | Lower | Upper |
|---|---|---|---|---|
| optoFGFR | RAF709 | 8.0643 | 7.2915 | 8.8371 |
| | U0126 | 12.8672 | 11.9596 | 13.7749 |
| | SCH772984 | 1.3799 | 1.2593 | 1.5005 |
| optoFGFR + 100 µM SL0101 | RAF709 | 3.5101 | 3.0769 | 3.9433 |
| | U0126 | 5.7922 | 5.2616 | 6.3228 |
| | SCH772984 | 0.4016 | 0.3363 | 0.4669 |

**Appendix Table S7: $EC_{50}$ with upper and lower values for the fit shown in Figure EV5E**

#### Appendix Movies

**Appendix Movie S1: ERK dynamics in response to a transient optoFGFR input.** Cells stably expressing ERK-KTR-mRuby2, H2B-miRFP703 and optoFGFR-mCitrine were stimulated with a 470 nm light pulse (18 mJ/cm$^2$) at t = 9 minutes (blue top band). ERK-KTR and H2B were acquired at 1-minute intervals with a 20x air objective. OptoFGFR was acquired at the end of the experiment (t = 40 minutes). ERK-KTR nuclear signal was segmented based on the H2B nuclear marker (green circle). ERK-KTR cytosolic signal was extracted in a 4-pixels ring around the nucleus. The ring was obtained by expanding the nuclear mask by 2-pixels (blue circle) to exclude the blurred edges of the nucleus and further expanding that new mask by 4 pixels in a threshold-based manner (pink circle). Single-cell ERK activity was then calculated as the cytosolic/nuclear ERK-KTR ratio. Scale bar: 50 µm.

**Appendix Movie S2: OptoFGFR dimerization dynamics.** Cell stably expressing optoFGFR-mScarlet was imaged at 20-second intervals with a 100x TIRF objective and stimulated with a 470 nm light pulse at t = 300 seconds (blue top band). The Ilastik pixel classification module was used to manually annotate and segment optoFGFR dimers/oligomers events without quantifying the endocytic vesicles (see Appendix Figure S1B). OptoFGFR dimerization was then quantified by computing the mean of pixel intensities from the binarized mask obtained with Ilastik using Fiji. Scale bar: 5 µm.

**Appendix Movie S3: Different optoFGFR inputs trigger transient, oscillatory and sustained ERK dynamics.** Cells stably expressing ERK-KTR-mRuby2, H2B-miRFP703 and optoFGFR-mCitrine were stimulated with 470 nm light pulse (18 mJ/cm$^2$) at 2-minute intervals (blue top bands). ERK-KTR and H2B were acquired at 1-minute interval with a 20x air objective. OptoFGFR was acquired at the end of the experiment (t = 55 minutes). High optoFGFR level (cells 3, 4, and 5) lead to a sustained ERK activity, while low optoFGFR level (cells 1 and 2) lead to oscillatory ERK dynamics. Scale bar : 25 µm.
